# Nanoscopic dopamine transporter distribution and conformation are inversely regulated by excitatory drive and D2-autoreceptor activity

**DOI:** 10.1101/2021.03.09.434538

**Authors:** Matthew D. Lycas, Aske L. Ejdrup, Andreas T. Sørensen, Nicolai O. Haahr, Søren H. Jørgensen, Daryl A. Guthrie, Jonatan F. Støier, Christian Werner, Amy Hauck Newman, Markus Sauer, Freja Herborg, Ulrik Gether

## Abstract

The nanoscopic organization and regulation of individual molecular components in presynaptic varicosities of neurons releasing modulatory volume neurotransmitters like dopamine (DA) remain largely elusive. Here we show by application of several super-resolution microscopy techniques to cultured neurons and mouse striatal slices, that the dopamine transporter (DAT), a key protein in varicosities of dopaminergic neurons, exists in the membrane in dynamic equilibrium between an inward-facing nanodomain-localized and outward-facing unclustered configuration. The balance between these configurations is inversely regulated by excitatory drive and by DA D2-autoreceptor activation in manner dependent on Ca^2+^-influx via N-type voltage-gated Ca^2+^-channels. The DAT nanodomains contain tens of transporters molecules and overlap with nanodomains of PIP2 (phosphatidylinositol-4,5-bisphosphate) but show little overlap with D2-autoreceptor, syntaxin-1 and clathrin nanodomains. Summarized, the data reveal a mechanism for rapid alterations in nanoscopic DAT distribution and show a striking link of this to the conformational state of the transporter.

## INTRODUCTION

In its 15 years of utilization, super-resolution microscopy has improved our insights into the molecular organization and dynamic regulation of pre- and postsynaptic structures in the brain. Indeed, it has become clear that regulatory processes of fundamental importance for spatial and temporal control of synaptic signaling occur at a nanoscale level. In excitatory synapses, for example, it is now believed that AMPA-type and NMDA-type ionotropic glutamate receptors segregate into distinct nanodomains in the postsynaptic density and that this co-organization plays a critical role in synaptic physiology (Choquet and Hosy, 2020; Groc and Choquet, 2020). Such studies highlight the importance of observing past the diffraction limit of visible light while still leveraging labeling strategies and live imaging compatibility that light microscopy provides (Choquet and Hosy, 2020; Groc and Choquet, 2020; Liu and Kaeser, 2019).

Much less is known about the nanoscale architecture of the presynaptic compartment in neurons releasing modulatory transmitters like dopamine (DA). DA neurons project from the midbrain to the basal ganglia, the limbic system and the prefrontal cortex to mediate the role of DA in motor functions, reward mechanisms and learning processes (Bjorklund and Dunnett, 2007; Iversen, 1975; Tritsch and Sabatini, 2012). The DA neurons have remarkable axonal arbors with numerous release sites (i.e. varicosities) (Descarries et al., 1996; Giguere et al., 2019), and DA neurotransmission differs from classical fast synaptic transmission by operating perhaps primarily via “volume transmission”; that is, DA is predominantly released from non-synaptic release sites to act on target cells often located micrometers away (Borroto-Escuela et al., 2018; Descarries *et al*., 1996). It remains nonetheless elusive how the different molecular components in the dopaminergic release sites need to be organized at a nanoscale level and individually regulated to enable proper neurotransmitter release and reuptake.

A key regulator of DA neurotransmission, the dopamine transporter (DAT), is highly expressed in the varicosities of DA neurons and removes released DA from the extracellular space. DAT has received attention as the primary target for therapeutics, such as methylphenidate, and drugs of abuse such as methamphetamine and cocaine (German et al., 2015; Kristensen et al., 2011; Torres and Amara, 2007). Not surprisingly, DAT is subject to tight regulation through several mechanisms involving post-translational modifications, protein-protein interactions and protein-lipid interactions (Bermingham and Blakely, 2016; Eriksen et al., 2010; German *et al*., 2015; Kristensen *et al*., 2011; Ramamoorthy et al., 2011; Torres, 2006). Moreover, we recently showed by application of super-resolution imaging that DAT regulation might involve mechanisms that cannot be revealed by more classical methods (Rahbek-Clemmensen et al., 2017). Our data showed that DAT localizes to discrete nanodomains in the plasma membrane of cultured DA neurons and that activation of NMDA-receptors reduced DAT nanodomain localization (Rahbek-Clemmensen *et al*., 2017). Although this indicated that nanoscale dynamics in dopamine varicosities could be critical for controlling DA neurotransmission, the mechanisms and functional implications remain unknown.

Here we use several super-resolution imaging techniques, including Photoactivated Localization Microscopy (PALM) (Betzig et al., 2006), direct Stochastic Optical Reconstruction Microscopy (dSTORM) (Heilemann et al., 2008; Huang et al., 2009), 10 x Expansion Microscopy (ExM) (Chen et al., 2015; Tillberg et al., 2016; Truckenbrodt et al., 2019) and single-particle tracking dSTORM (spt-dSTORM) (Manley et al., 2008) together with new fluorescent cocaine-analogues (Guthrie et al., 2020) and new data analysis tools, to investigate how DAT and associated molecular components at DA varicosities are dynamically regulated. In summary, our experimental efforts demonstrate that excitatory drive and D2-autoreceptor activity inversely regulate in a Ca^2+^ dependent manner the conformational state of DAT and its localization in DA varicosities to PIP2 enriched nanodomains separate from syntaxin-1 (STX-1), D2-autoreceptor or clathrin nanodomains. In this way, our study uncovers a hitherto unknown level of regulation of an essential membrane protein in a modulatory transmitter terminal.

## RESULTS

### DAT shows nanodomain distribution in cultured neurons and striatal slices

There are a variety of strategies to resolve biological structures past the diffraction limit of light, but each method is associated with strengths, compromises, and potential artefacts. True nanoscopic organizational principles should therefore be observed across a variety of super-resolution methods as well as model systems. Our previous dSTORM work suggested that DAT distributes into nanodomains at DA varicosities of cultured midbrain DA neurons (Rahbek-Clemmensen *et al*., 2017). To substantiate this finding, we decided to assess DAT distribution across a variety of super-resolution methods and ultimately investigate whether DAT is distributed into nanodomains in intact tissue. First, by magneto-transfection we expressed in the DA neurons DAT with the photoconvertible fluorescent protein mEOS2 (Baker et al., 2010) fused to the N-terminus. This enabled us to visualize distribution of the transporter without use of antibodies. Importantly, images of tyrosine hydroxylase (TH) positive varicosities revealed a clustered distribution of the mEOS2-DAT signal surrounding the cytosolic TH signal (Figure 1A-D), similar to that seen by imaging the endogenous DAT by dSTORM (Rahbek-Clemmensen *et al*., 2017). We next employed astigmatic 3D-dSTORM to show that the DAT nanodomains were localized to the periphery of the varicosities, substantiating that DAT is found mainly in the plasma membrane (Figure 1E-H). To assess the distribution of DAT in intact tissue, we analyzed coronal mouse brain slices (Figure 1I-L). Widefield imaging exposed the DAT positive varicosities and by dSTORM we could visualize DAT in single varicosities, showing a nanodomain distribution like that seen in cultured neurons (Figure 1I-L). To exclude that the observed distribution of DAT was not caused by artifacts introduced by the single molecule localization microscopy approach (e.g., “overcounting” (Shivanandan et al., 2014)), we turned to ExM where nanoscale spatial resolution is obtained by expanding the tissue sample using a polymer system (Chen *et al*., 2015). We employed a 10x expansion protocol (Truckenbrodt *et al*., 2019) to mouse striatal slices before imaging DAT in the slices by use of a spinning disk confocal microscope (Figure 1M-P). In the stained expanded sample, we captured the intermingled axonal network of DA neurons while at the same time observing clustering of DAT into nanodomains in varicosities (Figure 1M-P). Summarized, the distribution of DAT into nanodomains in the plasma membrane of DA neurons was substantiated by several high-resolution imaging techniques.

**Figure 1.**
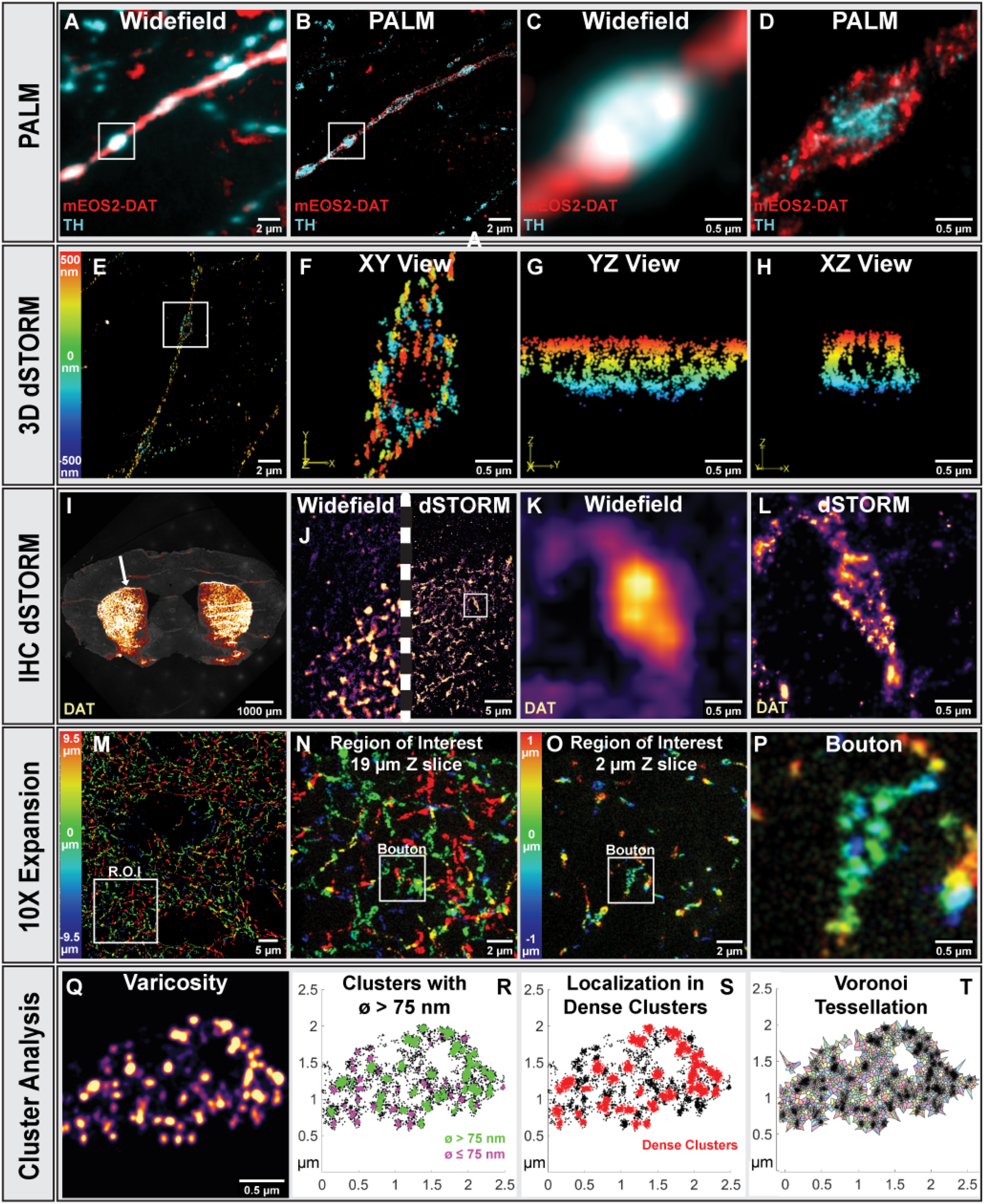
DAT shows nanodomain distribution in cultured neurons and striatal slices. (A-D) Visualization of mEOS2-DAT overexpressed in cultured DA neurons. Endogenous TH was labeled to identify DA neurons. (A, C) Widefield images (C is boxed region in A). (B, D) PALM/dSTORM images (D is boxed region in B). (E-H) 3D dSTORM image of DAT in DA neurons (color-coded based on depth). (E) Neuronal extension. (F) Varicosity (boxed region in E) (XY perspective). (G) YZ perspective. (H) XZ perspective. (I-L) Mouse brain coronal section immunolabeled for DAT. (I) brain slice with striatal DAT staining. (J) Comparison of widefield with dSTORM. (K) Widefield view of varicosity. (L) dSTORM image. (M-P) 10 x Expansion Microscopy (ExM) on mouse striatal slice immunolabeled for DAT. N shows region of interest (R.O.I.) in M with the full 19 µM Z slice (color-coded based on depth); O is R.O.I. in M with just 2 µM of the Z axis. (P) Single varicosity identified in O. (Q-T) Cluster metrics demonstrated on a varicosity immunolabeled for DAT. Q, dSTORM of single varicosity. (R, S) Identification of clusters by DBSCAN; clusters with diameter >75 nm in green, clusters <75 nm in magenta; dense clusters (>80 localizations within a radius of 50 nm) in red. (T) DAT localizations segmented by Voronoi tessellation. See also Figures S1 and S2.

Having reaffirmed the clustered organization of DAT with different techniques and in intact brain tissue, we decided to focus on the dSTORM technology in our subsequent studies because of its quantitative nature and single-molecule sensitivity. To derive quantitative measures for the clustered distribution of DAT, we employed the DBSCAN algorithm (density based spatial clustering of applications with noise) (Ester et al., 1996; Rahbek-Clemmensen *et al*., 2017) and extracted two key parameters from the analysis: the size of the nanoclusters and the density of localizations within the clusters (Fig. 1Q-S). The size was quantified as the fraction of clusters with a diameter >75 nm (Figure 1R). This size was chosen as threshold to exclude any clusters that might be the result of amplified labeling of single proteins and thus might arise from the antibody labeling methodology. By assessing the density of localizations as well, we could determine changes in the number of molecules in a cluster without a change in cluster size. Dense clusters were defined as clusters with >80 localizations within a radius of 50 nm (dense clusters in red) (Figure 1S). In selected cases, we also employed Voronoi tessellation to compare the density distributions of localization following pharmacological treatments in a parameter free fashion (Levet et al., 2019). In this use of Voronoi tessellation, clusters themselves were not identified but localization density changes could be detected with greater sensitivity (Figure 1T).

To estimate how many DAT molecules that are present in an average DAT nanodomain defined by the DBSCAN analysis, we used a protocol modified from (Ehmann et al., 2014; Siddig et al., 2020) and labeled cultured DA neurons with sequential dilutions of the primary DAT antibody. This enabled us to define that a minimum cluster is made from ∼three localizations and that an average large cluster contains ∼33 copies of the antibody corresponding to 50-100 DAT molecules as a rough estimate (Figure S1). The same data series lent itself perfectly for cluster analysis by varied label density cluster verification, enabling us to distinguish between random clustering arising from multiple observations of single fluorophores and true clustering (Baumgart et al., 2016). In agreement with our data in DAT expressing Cath.a-differentiated (CAD) cells (Rahbek-Clemmensen *et al*., 2017), our analysis supported that the clustering behavior of DAT in cultured neurons reflect true nanocluster formation (Figure S2).

### Nanodomain localized DAT is preferentially in an inward-facing conformation

To assess putative functional implications of the nanoclustered distribution of DAT, we investigated whether this could involve a conformational bias with nanoclustered DAT having a different conformation than unclustered DAT. Cocaine and cocaine-like inhibitors are known to bind with higher affinity to DAT in the outward-facing conformation than in the inward-facing conformation (Beuming et al., 2008; Loland et al., 2008; Newman et al., 2019). We envisioned accordingly (and in line with (Lebowitz et al., 2019)) that an indirect measure of the conformational state of DAT inside and outside the nanodomains could be obtained by parallel imaging of binding to DAT of a fluorescently tagged cocaine-analogue and of DAT itself; that is, as a result of its conformational bias the cocaine-analogue would be expected to bind more to outward-facing than to inward-facing transporters at a subsaturating concentration. We labeled cultured DA neurons and mEOS2-DAT expressing CAD cells with DG3-63 (10 nM), an analogue of cocaine tagged with JF_646_ suited for dSTORM (Figure 2A-D). The DG3-63 signal generally appeared somewhat less clustered as compared to that observed by our other approaches (see Figure 1). This might indicate a preferential binding of DG3-63 to unclustered DAT, which was supported by dual-color images of DG3-63 labelled DAT and mEOS-DAT, showing a more dispersed DG3-63 signal as compared to a more clustered mEOS2-DAT signal (Figure 2E-G). Importantly, the DG3-63 signal was specific as it was essentially eliminated by excess of unlabeled cocaine (Figure 2H). To quantify whether DG3-63 preferentially bound to unclustered DAT, we used a modification of the principles outlined by the colocalization extension of the Voronoi tessellation algorithm (Levet *et al*., 2019) where associated DG3-63 was identified by including DG3-63 localizations within 25 nm from an mEOS2-DAT localization (Figure 2I-K). The total mEOS2-DAT Voronoi tessellated area distribution showed a preference for smaller tessellated areas consistent with a clustered distribution (Figure 2L). In contrast, the mEOS2-DAT-associated DG3-63 localizations resided on larger Voronoi tessellated areas; hence, DG3-63 binding was not enriched in the DAT nanodomains containing the highest density of DAT molecules (Figure 2L). This suggests that nanoclustered DAT are less prone to bind DG3-63, consistent with a more inward-facing transporter conformation or a less accessible binding site. Interestingly, by analyzing the distribution of mEOS2-DAT after treatment with an excess, and thereby saturating concentration, of cocaine, we observed a dispersal of the signal. This supports that forcing the transporter into an outward-facing conformation may release the transporter from the nanodomains (Figure S3A).

**Figure 2.**
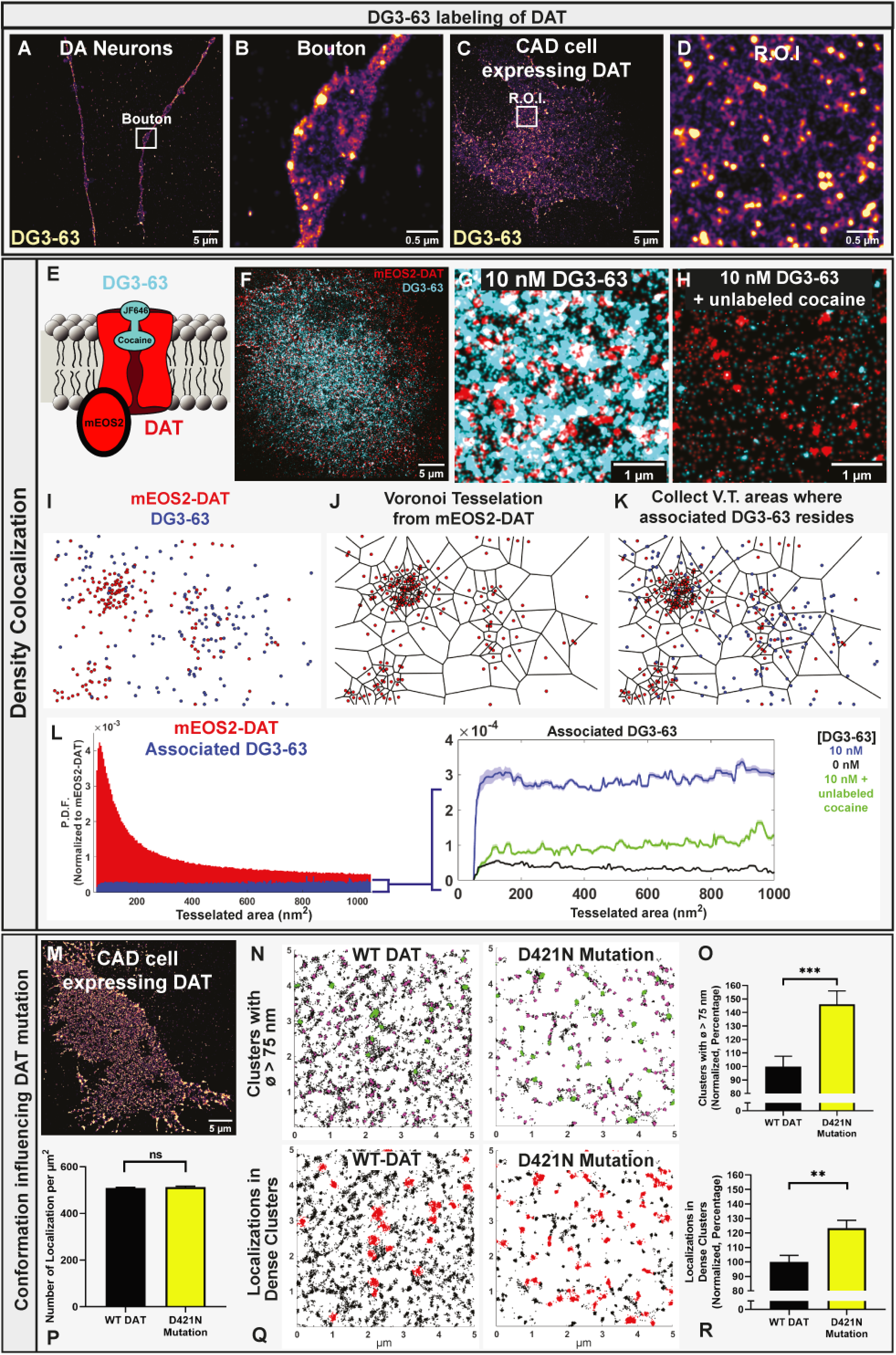
Nanodomain localized DAT is preferentially in an inward-facing conformation. (A-D) dSTORM on DA neurons and DAT expressing CAD cells labeled with the fluorescent cocaine analogue DG3-63 (10 nM). (A) Axonal extension. (B) varicosity (box in A). (C) CAD cell. (D) R.O.I. from C. (E-H) The density-based association of DG3-63 to mEOS2-DAT. (E) Cartoon of labeling strategy. (F) dSTORM/PALM image of CAD cell expressing mEOS2-DAT (red) and labeled with DG3-63 (10 nM)(blue). (G) Close-up dSTORM/PALM image of CAD cell expressing mEOS2-DAT (red) and labeled with DG3-63 (10 nM) (blue). (H) The DG3-63 signal is blocked by 100 µM cocaine. (I-K) Voronoi tessellation based colocalization algorithm. Tessellation is performed on the mEOS2-DAT data and areas are marked for when the mEOS2-DAT localization has a DG3-63 localization within 25 nm distance. (L) Left, Total mEOS2-DAT Voronoi tesselated area distribution (red) (probability density function (P.D.F.) and tessellated areas of mEOS2-DAT associated with DG3-63 (blue). Right, tessellated areas for mEOS2-DAT associated with DG3-63 (blue) compared to that associated in presence of saturating cocaine. (M-O) TIRF/dSTORM of WT DAT and inward-facing mutant (D421N). (M) Example image of DAT expressed in CAD cells. (N) Example images of clusters with diameter >75 nm (green) in WT DAT and D421N Dcells. (O) Normalized fraction of clusters with diameter >75 nm (in %), means ± S.E., from 38 WT DAT cells and 42 D421N cells, 3 transfections, ***p<0.001, unpaired t-test. (P-O) dSTORM of WT DAT and inward-facing mutant (D421N). (P) Number of localizations per µm^2^ for WT DAT and D421N, means ± S.E., unpaired t test, p= 0.4. (Q) Example image of dense clusters (red) of WT DAT and D421N (radius 100 nm, >100 localizations). (R) Fraction of localizations in dense clusters (in % of WT DAT), means ± S.E., from 38 WT DAT cells and 42 D421N cells, 3 transfections, **p<0.01, unpaired t-test. See also Figure S3.

To further address this question, we examined the effect of the DAT inhibitors nomifensine, JHW007 and ibogaine on DAT nanoclustering in CAD cells transiently expressing WT DAT. We decided to use dSTORM and thus immunolabeling rather than mEOS2-DAT to make the data more compatible with our analyses on neurons and brain slices. Of the different inhibitors, nomifensine is a cocaine-like inhibitor prone to self-administration (Spyraki and Fibiger, 1981) a and thus prone to stabilize an outward-facing conformation (Beuming *et al*., 2008; Loland *et al*., 2008; Newman *et al*., 2019). JHW007 and ibogaine, however, are known to promote an inward-facing conformation of the transporter (Beuming *et al*., 2008; Kasture et al., 2016; Loland *et al*., 2008; Newman *et al*., 2019). Importantly, a Voronoi tessellation analysis of the data showed that while nomifensine decreased nanoclustering, both JHW007 and ibogaine increased DAT clustering, consistent with the notion that an inward-facing conformation promotes nanodomain localization (Figure S3B, C).

We further rationalized that if the nanoscale DAT distribution reflects its molecular conformation, then a redistribution would be expected for a DAT variant with a conformational bias. Accordingly, we compared in transfected CAD cells clustering of WT DAT with that of a disease-associated DAT mutation (D421N) known to be shifted towards an inward-facing conformation (Hansen et al., 2014; Herborg et al., 2018) (Figure 2M-R). In agreement with the idea that inward-facing DAT will display preferential localization to nanodomains, analysis of the resulting dSTORM data revealed for D421N compared to WT both a higher fraction of large clusters (diameter >75 nm) and a higher fraction of localizations in dense clusters (Figure 2N, O, Q, R) without a change in total number of detected localizations (Fig. 2P). Summarized, the data support that DAT preferentially assumes an inward-facing conformation in DAT nanodomains while the conformation outside the nanodomains conceivably is more outward-facing.

### NMDA receptor activation declusters DAT in striatal slices

We next assessed whether DAT nanodomain distribution was sensitive to NMDA stimulation in intact tissue (Figure 2I-P) as we previously observed in cultured neurons (Rahbek-Clemmensen *et al*., 2017). Notably, NMDA receptors are present in DA terminals where they can stimulate DA release (Salamone et al., 2014). Acute coronal striatal slices were treated for 5 min with vehicle, NMDA or NMDA plus the NMDA receptor antagonist (2R)-amino-5-phosphonovaleric acid (AP5) before DAT immunolabeling and dSTORM (Figure 3A). Because each dSTORM image contained a vast number of DAT positive varicosities, we developed a new collection algorithm for automated and unbiased isolation of these varicosities, allowing subsequent quantification of DAT clustering (Figure 3B, C). Following NMDA treatment, both the fraction of clusters >75 nm and the faction of localizations in dense clusters decreased; an effect that was muted by the NMDA antagonist AP5 (Figure 3D, E). Note that the comparisons were normalized to the control sample from the individual imaging session to correct for unavoidable variability in sample preparations and between imaging sessions. Importantly, the data were further supported by Voronoi tessellation showing that NMDA, but not NMDA + AP5, reduced the fraction of smaller tessellated areas and thus clustering (Figure 3F). We conclude that local activation of NMDA receptors in intact tissue promotes dispersal of DAT nanodomains.

**Figure 3.**
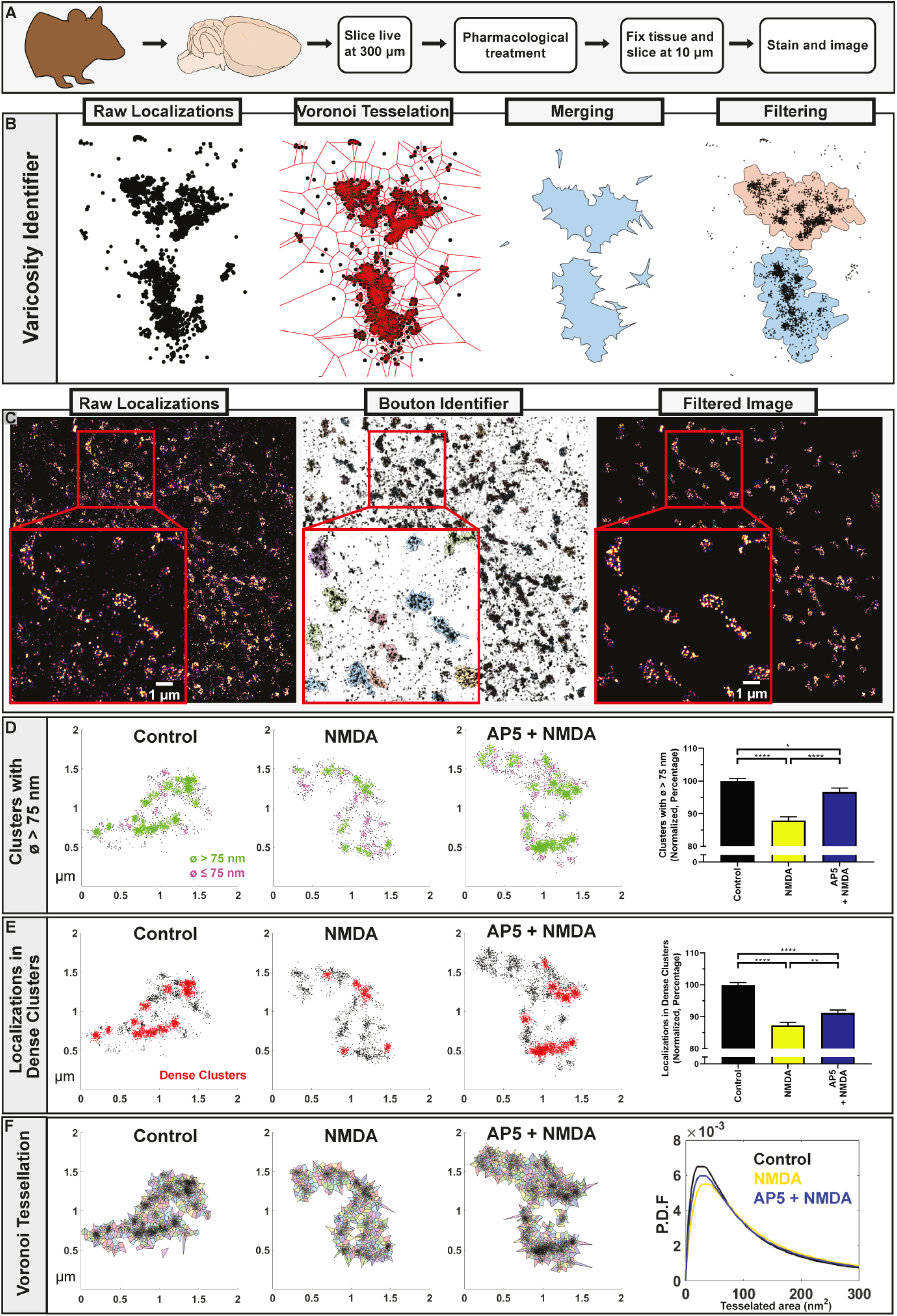
NMDA receptor activation declusters DAT in mouse brain slices. (A) Workflow diagram. (B) Data processing steps to isolate single varicosities from dSTORM images (see Methods). (C) Representative DAT dSTORM image before, during, and after varicosity identification. (D) Example varicosity from control, NMDA, and AP5 + NMDA treated slices (clusters with diameter >75 nm in green, clusters <75 nm in magenta). Right, Normalized fraction of DAT clusters >75 nm (in %), means ± S.E., from 3 experiments with analysis of 1793 control, 1102 NMDA and 1033 NMDA + AP5 varicosities; one-way ANOVA, ****p<0.0001, *p<0.05. (E) Example varicosity from control, NMDA (20 µM), and AP5 + NMDA (20 µM) treated slices (dense clusters in red, >80 localizations, radius 50 nm). Right, Normalized fraction of DAT localizations in dense clusters (in %), means ± S.E.; one-way ANOVA, ****p<0.0001, ***p<0.001, **p<0.01. (F) The probability density functions (P.D.F.) for the Voronoi tessellated areas averaged by varicosity.

### DAT nanodomain localization is regulated by neuronal activity

We speculated whether persistent changes in membrane potential and thereby in neuronal excitability would affect DAT nanodomain distribution. To test this, we expressed the bacterial sodium channel (mNaChBac) or the inward-rectifier potassium channel (Kir2.1) (Lin et al., 2010; Xue et al., 2014) in cultured DA neurons to make them more or less excitable, respectively (Lin *et al*., 2010). As a control, we also expressed an inactive mutant of mNaChBac (mNaChBac MUT) (Lin *et al*., 2010; Xue *et al*., 2014). In agreement with selective expression in DA neurons, we confirmed that transduced cells were also DAT positive (Figure S4). Electrophysiological recordings on single neurons showed that the pacemaker firing activity of DA neurons (Rayport et al., 1992) was enhanced in mNaChBac-expressing neurons and reduced in Kir2.1 expressing neurons with no significant change in mNaChBac MUT-expressing neurons (Fig S4). Analysis by dSTORM of DAT distribution in mNaChBac expressing neurons showed decreased clustering, while for Kir2.1 expressing neurons we saw a trend towards an increased fraction of large clusters and a significant increase in localizations within large dense clusters (Fig. 4A-D). No significant effect was seen for mNaChBac MUT (Figure 4C, D). Thus, DAT nanodomain localization is in cultured DA neurons regulated by persistent changes in excitability or firing activity with increased activity leading to dispersal of DAT from nanodomains and decreased activity leading to increased nanodomains localization.

**Figure 4.**
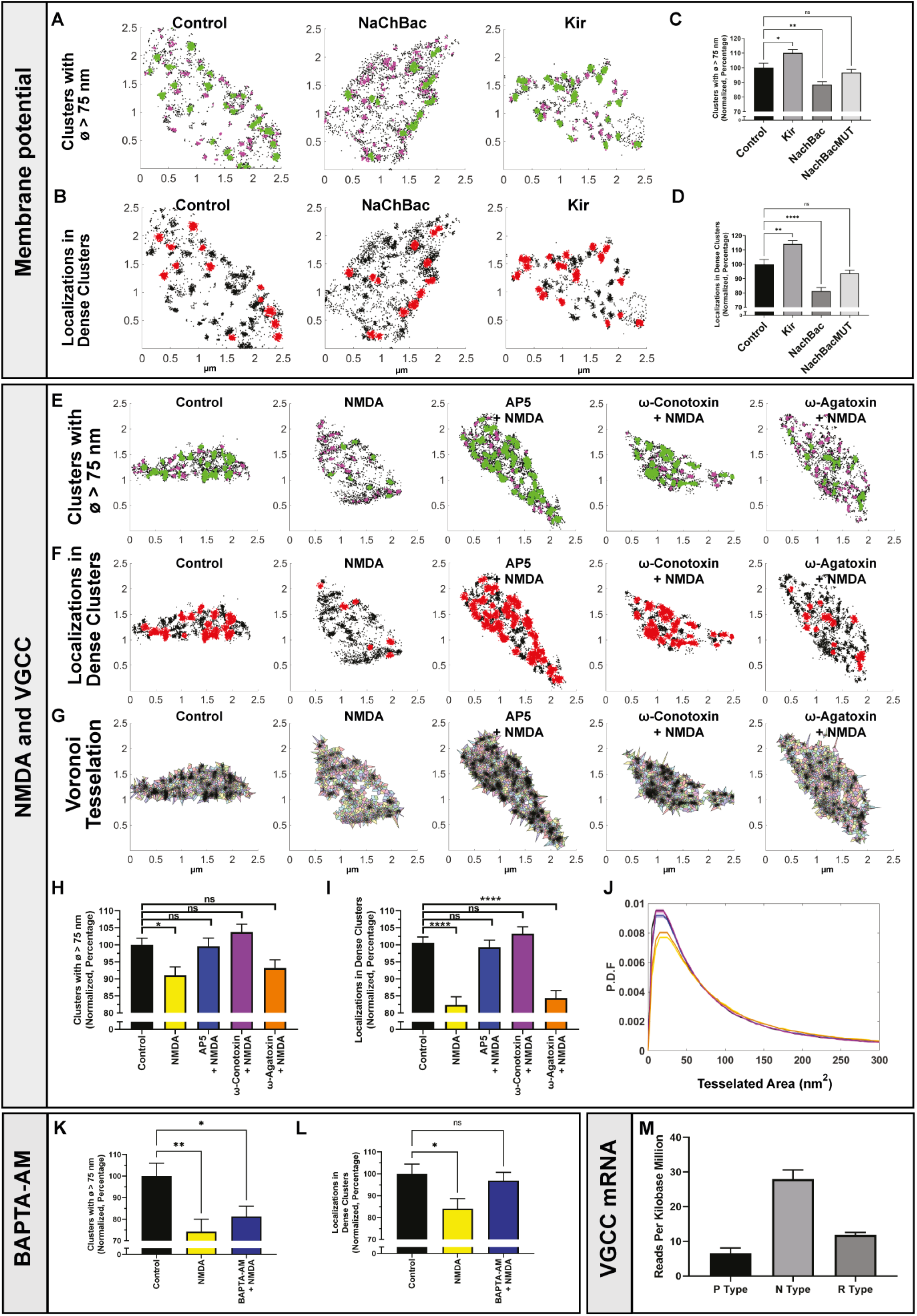
DAT nanodomain localization is regulated by neuronal activity in a Ca^2+^ dependent manner. (A-D) Expression of the Na^+^ channel mNaChBac or the K^+^ channel Kir2.1 in DA neurons. Example varicosities from DAT dSTORM of control, mNaChBac or Kir2.1 expressing DA neurons. (A) DAT clusters with diameter >75 nm in green, clusters <75 nm in magenta. (B) Dense DAT clusters in red (>80 localizations, radius 50 nm). (C) Normalized fraction of clusters >75 nm (in %). (D) Normalized fraction of DAT localizations in dense clusters (in %), means ± S.E.; data from 3 cultures and 128 control, 290 mNaChBac, 220 Kir2.1, and 211 mNaChBacMUT varicosities; one-way ANOVA, ****p<0.0001, **p<0.01, *p<0.05, n.s., not significant. (E-J) The effect of NMDA on DAT nanodomains is blocked by the N-type VGCC blocker ω-conotoxin. Example varicosities from dSTORM of DA neurons subject to 5 min treatment with vehicle, NMDA (20 µM), AP5 (100 µM) + NMDA (20 µM), ω-Conotoxin (1 µM) + NMDA (20 µM) or ω-Agatoxin (1 µM) + NMDA (20 µM). (E) DAT clusters with diameter >75 nm in green, clusters <75 nm in magenta. (F) Dense DAT clusters in red (>80 localizations, radius 50 nm). (G) localizations segmented by Voronoi tessellation. (H, I) Normalized fraction of DAT clusters >75 nm (in %) (H) and normalized fraction of DAT localizations in dense clusters (in %) (I), means ± S.E.; one-way ANOVA, ****p< <0.0001, *p<0.05, n.s., not significant. (J) Probability density functions (P.D.F.) (curves color-coded as in the bar diagrams) for the Voronoi tessellated areas averaged by varicosity. Data are from 3 cultures and 190 control, 147 NMDA, 170 AP5, 180 ω-Conotoxin and 154 ω-Agatoxin varicosities, (K, L) Pretreatment (30 in) with BAPTA-AM inhibits the dispersing effect of NMDA on DAT nanodomains. (K) Normalized fraction of DAT clusters >75 nm (in %) and (L) normalized fraction of DAT localizations in dense clusters (in %) for control, NMDA (20 µM) and NMDA (20 µM) + BAPTA-AM (25 µM), means ± S.E.; data from 3 cultures and includes analysis of 66 control, 72 NMDA and 70 BAPTA-AM + NMDA varicosities, one-way ANOVA, **p<0.01, *p<0.05, n.s., not significant. (M) mRNA expression for P, R, and N-type VGCC in DA neurons based on single cell RNAseq data (see Methods). See also Figure S4 and Movies S1-2.

### DAT nanodomain localization is regulated by Ca^2+^

Because of the effect of neuronal excitability, we decided to investigate putative involvement of Ca^2+^ influx via voltage-gated Ca^2+^ channels (VGCC) in NMDA-induced DAT declustering. Note that both P/Q type and N-type VGCCs should be present in DA terminals (Turner et al., 1993). DA neurons were stimulated with NMDA for 5 min with or without the N-type VGCC blocker ω-conotoxin or the P/Q type VGCC blocker ω-agatoxin. While we observed no effect of ω-agatoxin, the dispersing effect of NMDA on DAT clustering was essentially abolished by ω-conotoxin. Similarly, Voronoi tessellation analysis showed that ω-conotoxin, but not ω-agatoxin, blocked the NMDA-induced shift away from small, tessellated areas and thus blocked the dispersal of DAT nanodomains (Figure 4E-J, Movies S1-2). We next asked whether Ca^2+^ would be important *per se* by preincubating the neurons with the cell permeable Ca^2+^-chelator BAPTA-AM. Indeed, BAPTA-AM blunted the response to NMDA (Figure 4K-L). Of note, the pharmacological data fitted well with available single-cell mRNA sequencing data on DA neurons supporting highest transcription of N-type VGCC in DA neurons (Figure 4M) (see Methods for data bases used). Taken together, NMDA-induced dispersal of DAT nanodomains appears to occur via Ca^2+^-dependent mechanism involving the activity of N-type VGCCs.

### DAT nanodomain localization is regulated by the D2 autoreceptor

D2-autoreceptors in DA neurons act as negative feedback regulators decreasing neuronal excitability and firing activity by promoting opening of inhibitory G-protein-activated inwardly rectifying potassium channels (Beckstead et al., 2004; Kuzhikandathil et al., 1998). We surmised that D2-autoreceptor activity might regulate localization of DAT to nanodomains and tested this by determining the effect of the D2 receptor antagonist haloperidol and the D2 receptor agonist quinpirole. Interestingly, incubation with 10 nM haloperidol for 5 min resulted in dispersal of DAT nanodomains with a reduced fraction of clusters >75 m, fewer localizations in dense clusters and a reduced fraction of smaller Voronoi tessellated areas (Figure 5A-F, Movies S1, S3). This effect is likely the result of haloperidol blocking the effect on the D2-autoreceptors of DA released in the culture as a consequence of DA neuronal pacemaker activity as shown in Figure S4. Also, co-incubation with an excess of quinpirole blocked the effect of haloperidol (Figure 5A-F). Moreover, the effect of haloperidol was accompanied by an increase in cytoplasmic free Ca^2+^ consistent with D2-autoreceptor activity suppressing neuronal firing (Figure S5A, B). This prompted us to test if the effect of haloperidol, like the effect of NMDA, was dependent on Ca^2+^-influx via N-type VGCCs. This appeared to be the case as ω-conotoxin, but not ω-agatoxin, eliminated the effect of haloperidol (Figure 5A-F). Summarized, our data support that D2-autoreceptors are preoccupied with DA in the neuronal culture and that the activity of these receptors can regulate DAT nanodomain localization via a Ca^2+^-dependent mechanism.

**Figure 5.**
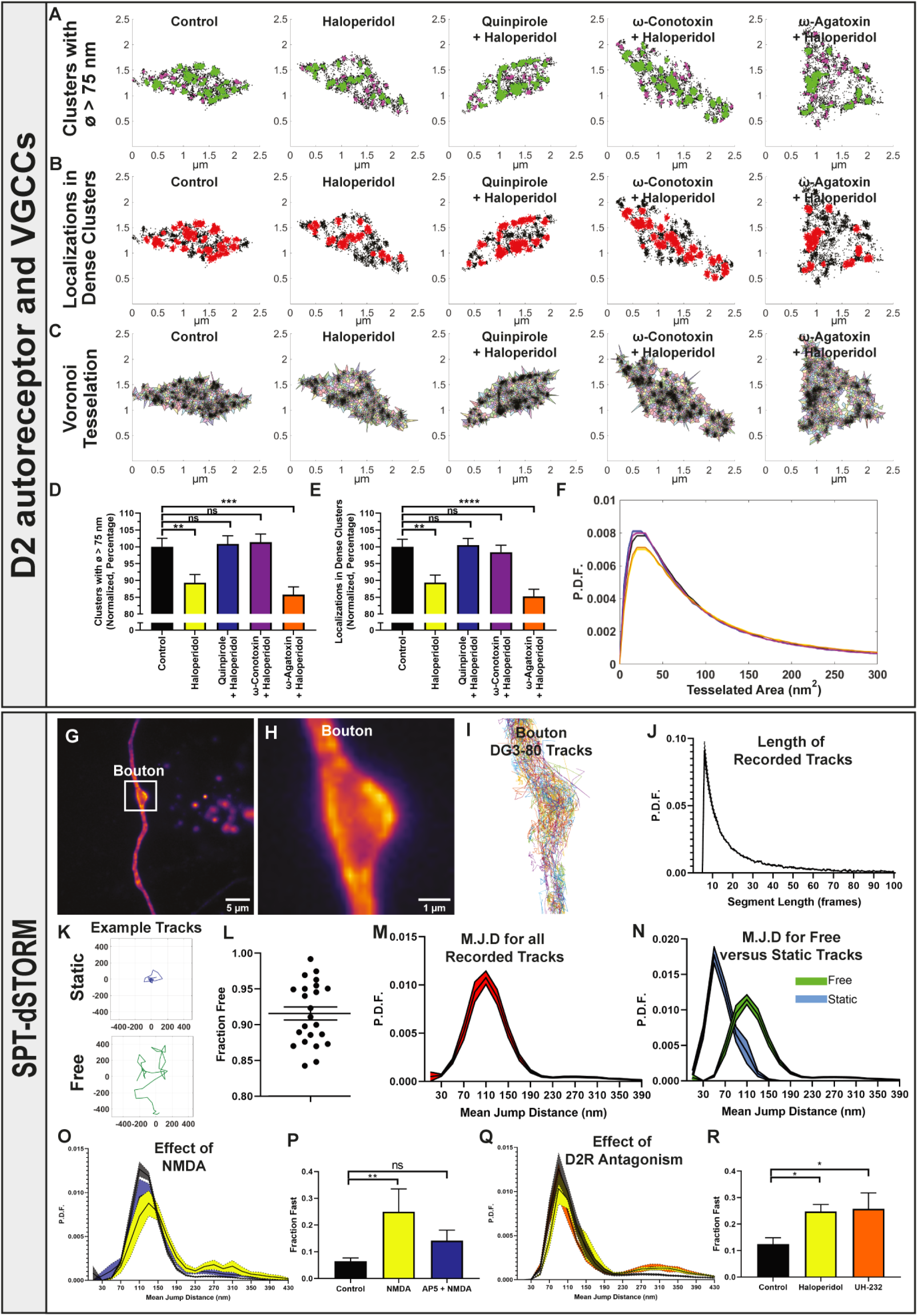
DAT nanodomain localization is regulated by the D2R and involves changes in lateral mobility. A-F) Dispersal of DAT nanodomains is promoted by the D2R antagonist haloperidol and blocked by the agonist quinpirole and by the N-type VGCC blocker ω-conotoxin. Example varicosities from dSTORM of DA neurons subject 5 min treatment with haloperidol (10 nM), haloperidol (10 nM) + quinpirole (50 µM), haloperidol (10 nM) + ω-conotoxin (1 µM) or haloperidol (10 nM) + ω-agatoxin (1 µM). (A) DAT clusters with diameter >75 nm in green, clusters <75 nm in magenta. (B) Dense DAT clusters in red (>80 localizations, radius 50 nm). (C) localizations segmented by Voronoi tessellation. (D, E) Normalized fraction of DAT clusters >75 nm (in %) (D) and normalized fraction of DAT localizations in dense clusters (in %) (E), means ± S.E.; one-way ANOVA, ****p<0.0001, ***p<0.001, **p<0.01, *p<0.05, n.s., not significant. (F) Probability density functions (P.D.F.) (curves color-coded as in the bar diagrams) for the Voronoi tessellated areas averaged by varicosity. Probability density functions (P.D.F.) (curves color-coded as in the bar diagrams) for the Voronoi tessellated areas averaged by varicosity. Data are from 3 cultures and 153 control, 166 haloperidol, 167 quinpirole, 178 ω-conotoxin and 176 ω-agatoxin varicosities, (G-R) Single particle tracking dSTORM (SPT-dSTORM) identifies changes in DAT mobility in response to NMDA and D2R antagonism. (G) Example live dSTORM images of DA neurons labeled with the fluorescent cocaine analogue, DG3-80. (H) Boxed region in B. (I) Detected tracks and (J) Distribution of segment lengths in varicosity from control SPT-dSTORM experiment (16 ms frame rate). (K) Example traces of a confined and an unconstrained freely moving particle (DG3-80 labeled DAT). (L) Fraction of unconstrained tracks out of the total number of tracks. Data are means of 23 tracks ± S.E. (M) Probability Density Function (P.D.F.) for total mean jump distances (M.J. D) (i.e. the distance an individual molecule/particle has traveled between each image) for control experiments. (N) P.D.F. when separated into static and freely moving particles. Among the freely moving particles, some moved very fast with M.J.D. of 200-350 nm. (O) NMDA (20 µM, 5 min) increases population of DAT molecules with M.J.D. > 200 nm, which is blunted by AP5 (100 µM). (P) Fraction of fast moving particles (> 200 nm), means ± S.E., data from 13 control, 4 NMDA and 5 NMDA+AP5 experiments; (Q, R) The D2R antagonists haloperidol (10 nM, 5 min) and UH-232 (1 µM) increase the population of DAT molecules with M.J.D. > 200 nm > 200 nm; M, Fraction of fast moving particles (> 200 nm), means ± S.E., data are from 10 control, 6 haloperidol and 5 UH-232 experiments; one-way ANOVA, *p<0.05; **p<0.01, n.s., not significant. See also Figure S5 and Movies S1, S3.

To further substantiate a correlation between D2-autoreceptor signaling, neuronal activity and DAT nanoclustering, we incubated cultured DA neurons with an antibody directed towards the luminal end of the presynaptic vesicular protein, synaptotagmin-1. Upon exocytotic release, this luminal epitope is exposed to the extracellular space and, thus, the degree of synaptotagmin-1 labeling can be considered a proxy for exocytotic release activity (Truckenbrodt et al., 2018). Indeed, when blocking the D2 autoreceptor with haloperidol, this did not only cause dispersal of DAT nanoclusters but was also accompanied by increased synaptotagmin-1 labeling as a likely result of increase exocytotic release activity (Figure S5C). Importantly, it is unlikely that the presumed increase in exocytotic release of DA directly regulates DAT and leads to dispersal of DAT nanodomains. Adding exogenous DA to the culture had no effect on DAT nanoclustering (Figure S3D) and, moreover, DAT nanoclustering was unaffected by noribogaine, while nomifensine caused a significant declustering of DAT (Figure S3D). This suggests that the transporter already is preoccupied by DA and thereby is active/cycling, which expectably pushes a fraction of the transporter molecules into an inward-facing nanoclustered configuration. In this scenario, no effect is expected of noribogaine as well as it is consistent with the declustering seen in response to nomifensine that will compete DA away from the transporter.

### Single particle tracking supports nanodomain dispersal in response to neuronal activity

To obtain a better understanding of the dynamics underlying DAT nanodomain localization, we employed SPT-dSTORM (Manley *et al*., 2008) to assess if dispersal of DAT from nanodomains is accompanied by increased lateral mobility of DAT in the membrane. To allow visualization in live cells, we labeled DAT in DA neurons with a Janelia Fluor®549-conjugated cocaine analogue, DG3-80, suited for live dSTORM (Guthrie *et al*., 2020) (Figure 5G, H). Individual DAT tracks were captured with lengths up to 1.6 sec (100 frames, 16 msec frame rate) with the majority being 10-20 frames (Figure 5I, J). Separating tracks into those reflecting confined movements and those reflecting freely moving particles showed that ∼90% of the tracks described freely moving particles (Figure 5K, L). Note that this distribution (with most of the labeled molecules being freely moving) likely reflects preferential binding of DG3-80 to unclustered DAT in the outward-facing conformation (Figure 2). The mean jump distance (i.e. the distance an individual molecule/particle has traveled between each image) was ∼110 nm for all the recorded tracks and ∼50 nm for static tracks only (Figure 5M, N). Interestingly, a jump distance of ∼110 nm is so fast that a transporter molecule might move 7-14 µm during a transport cycle estimated to last 1-2 sec (Prasad and Amara, 2001). Among the freely moving particles, there were even some that moved very fast with mean jump distance of 200-350 nm (Figure 5M-O). Both NMDA receptor activation and blockade of the D2-autoreceptors with either haloperidol or the neutral antagonist UH-232 increased the population of DAT molecules with mean jump distances >200 nm, indicative of a possible link between dispersal of DAT nanodomains and increased lateral mobility. Note that the similar effect of UH-232 and haloperidol strongly suggest that their effect reflects blockade of D2-autoreceptors activated by DA, which is released because of DA neuronal pacemaker activity, and not inhibition of constitutive receptor activity (Strange, 2008).

### DAT nanodomains overlap with PIP2 nanodomains

A key question is how the nanoscale distribution of DAT relates to other molecular components of DA terminals. We first turned to the plasma membrane phospholipid, PIP2, which can bind DAT and in doing so modulate DAT activity (Belovich et al., 2019). Dual-color dSTORM on PIP2 and DAT in DA neurons showed as expected (van den Bogaart et al., 2011), a distribution of the PIP2 signal into nanodomains (Figure 6). Strikingly, these nanodomains showed a remarkable overlap with DAT nanodomains with the Voronoi tessellation distributions revealing a large fraction of PIP2 localizations associated with dense DAT localizations (Figure 6A, B, F). As also seen for DAT domains, treatment with both NMDA and haloperidol reduced PIP2 localizations in large dense clusters while haloperidol also reduced cluster size. Importantly, all these changes were blocked by AP5 or quinpirole, respectively (Figure 6B-D). The NMDA and haloperidol-induced changes were reflected in the Voronoi tessellation distributions as well by showing dispersal of the PIP2 nanodomains and reduced PIP2 association to nanoclustered DAT (Figure 6E, F).

**Figure 6.**
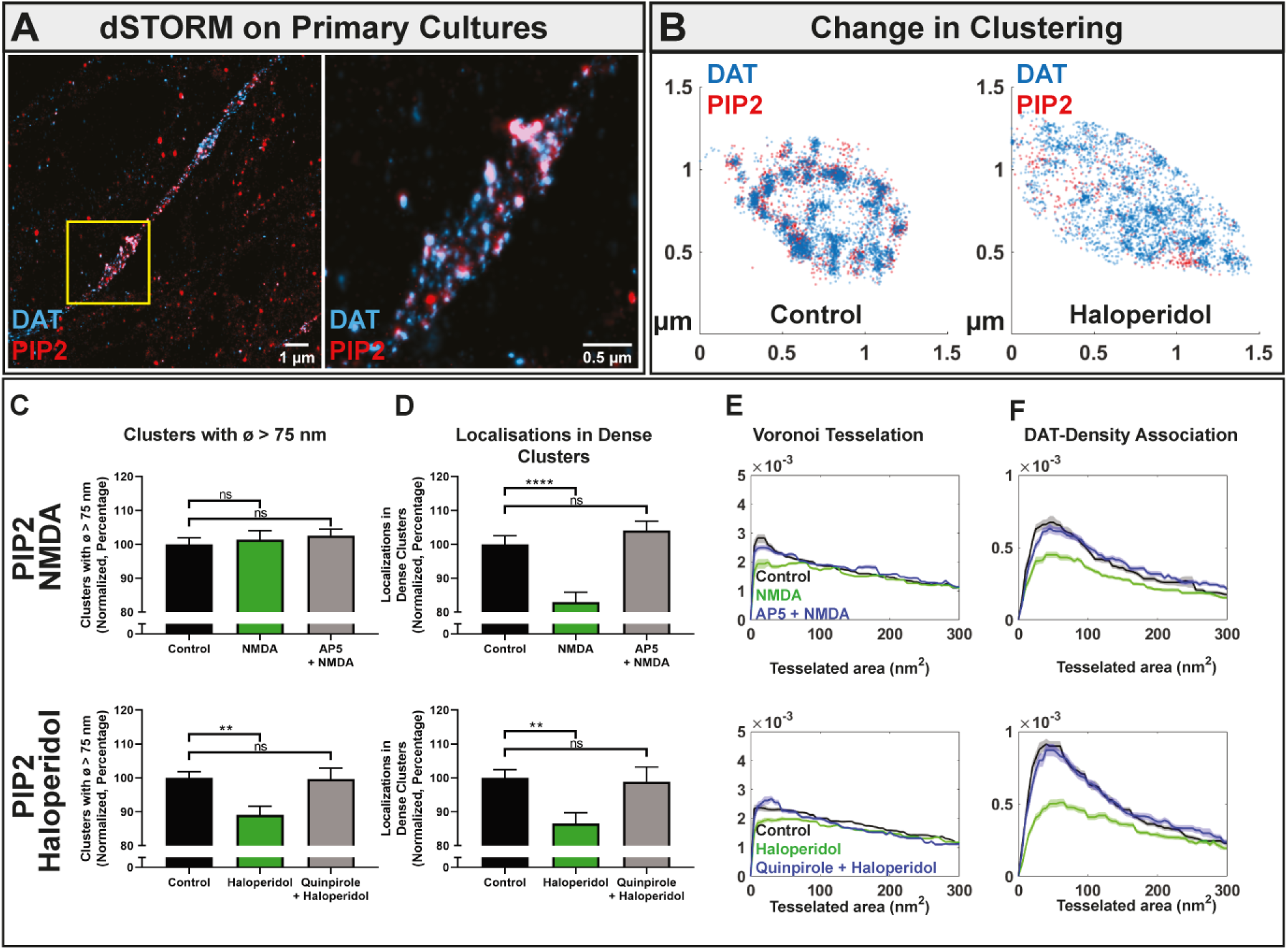
DAT nanodomains overlap with PIP2 nanodomains. (A) Example dual-color dSTORM images of phosphatidylinositol 4,5-bisphosphate (PIP2) (red, CF568) and DAT (blue, Alexa647) in a neuronal extension from cultured DA neurons (left) with close-up image of varicosity (right) corresponding to boxed region in the left image. (B) Example dual-color dSTORM images of varicosities from DA neurons showing localizations for PIP2 (red) and DAT (blue). Left, Control; Right, neurons exposed to haloperidol (10 nM, 5 min). (C-F) Effect of NMDA and haloperidol on PIP2 clustering. Upper panels, Control, NMDA (20 µM) and NMDA (20 µM) + AP5 (100 µM); Lower panels; Control, haloperidol (10 nM) and haloperidol (10 nM) + quinpirole (50 µM). (C) Normalized fraction of PIP2 localizations in large clusters (>75 nm in diameter) (in %) and (D), normalized fraction of PIP2 localizations in dense clusters (>80 localizations, radius 50 nm) (in %), means ±S.E., one-way ANOVA, ****p<0.0001, **p<0.01, n.s., not significant. (E) Probability density functions (P.D.F.) (color-coded as in the bar diagrams) for the Voronoi tessellated areas averaged by varicosity. (F) Probability density functions (P.D.F.) (color-coded as in the bar diagrams) of the association of PIP2 with DAT clusters determined by Voronoi tessellation-based association (see Figure 2) showing strong association PIP2 with DAT. Data are from 3 cultures and 137 control, 126 NMDA, 121 NMDA + AP5 varicosities; and 140 control, 102 haloperidol, 67 haloperidol + quinpirole varicosities,

### The D2R is also shows nanodomain distribution

We next analyzed the D2-autoreceptors that are found both in the somatodendritic compartment of DA neurons and in DA terminals. Due to known challenges with DA receptor antibodies, we both validated the specificity of the antibody in transfected HEK293 cells and on slices from D2 receptor knock-out mice (Figure S6). Subsequent dual-color dSTORM experiments on cultured DA neurons revealed a clustered distribution of the D2-autoreceptors with nanodomains of a similar size as those found for DAT (Figure 7A, B). Furthermore, like our findings for DAT and PIP2, NMDA and haloperidol promoted dispersal of the receptor domains (Figure 7B-E). Relatively few of the nanodomains, however, overlapped with the DAT domains as apparent from the Voronoi tessellation distributions (Figure 7A, B, F), and as for D2-autoreceptor association to DAT, we only observed marginal changes upon NMDA and haloperidol treatment (Figure 7F).

**Figure 7.**
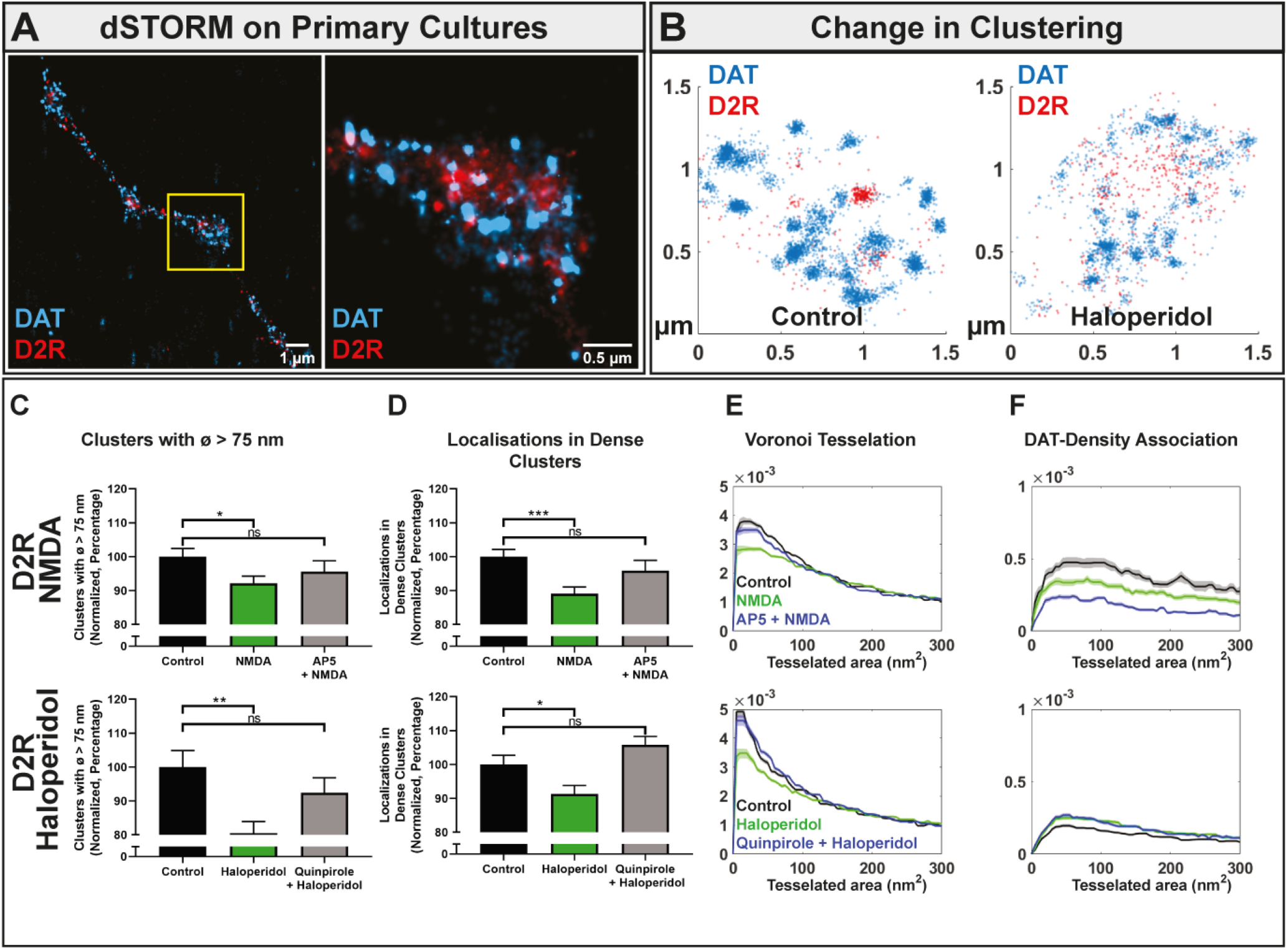
The D2R is also distributed into nanodomains. (A) Example dual-color dSTORM images of the D2R (red, CF568) and DAT (blue, Alexa647) in DA neuronal extension (left) with close-up image of varicosity (right) corresponding to boxed region in the left image. (B) Example dual color dSTORM images of varicosities from DA neurons showing localizations for D2R (red) and DAT (blue). Left, Control; Right, neurons exposed to haloperidol (10 nM, 5 min). (C-F) Effect of NMDA and haloperidol on D2R clustering. Upper panels, Control, NMDA (20 µM) and NMDA (20 µM) + AP5 (100 µM); Lower panels; Control, haloperidol (10 nM) and haloperidol (10 nM) + quinpirole (50 µM). (C) Normalized fraction of D2R localizations in large clusters (>75 nm in diameter) (in %) and (D), normalized fraction of D2R localizations in dense clusters (>80 localizations, radius 50 nm) (in %), means ±S.E., one-way ANOVA, ***p<0.001, **p<0.01, *p<0.05, n.s., not significant. (E) Probability density functions (P.D.F.) (color-coded as in the bar diagrams) for the Voronoi tessellated areas averaged by varicosity. (F) Probability density functions (P.D.F.) (color-coded as in the bar diagrams) of the association of D2R with DAT clusters determined by Voronoi tessellation-based association (see Figure 2). Data are from 3 cultures and data from 3 cultures and 112 control, 157 NMDA, 75 NMDA + AP5; and 179 control, 195 haloperidol and 150 quinpirole varicosities. See also Figure S5 and S6.

The findings prompted us to further assess the specificity of the pharmacological manipulations by imaging additional key membrane components in DA terminals including clathrin (Maxfield and McGraw, 2004) and the single transmembrane SNARE protein syntaxin1 (STX-1). Imaging of clathrin revealed a nanoclustered distribution, possibly corresponding to clathrin-coated pits (Li et al., 2018), that showed little co-localization with DAT and, importantly, essentially no change in distribution in response to NMDA (Figure S7). We also recapitulated nanodomain distribution of STX-1 that is known to exist in such domains in presynaptic terminals (Padmanabhan et al., 2020) as well as the protein has been proposed to bind DAT (Binda et al., 2008). However, the STX-1 domains showed little overlap with DAT and responded differently to NMDA with a decreased fraction of large clusters and an increased fraction of localizations in dense clusters (Figure S7). Note, however, that NMDA slightly increased localization of STX1 to unclustered DAT (i.e. the larger tessellated areas) according to the Voronoi tesselated distributions (Figure S6). In summary, several plasma membrane components of DA terminals show a nanoclustered distribution that, however, appears to be differentially regulated.

## DISCUSSION

DA neurons have remarkable axonal arbors with numerous presynaptic terminals or varicosities that serve as the principal architectural units responsible for release and reuptake of DA. Nonetheless, we still have a surprisingly poor understanding of the molecular organization of these varicosities, and what regulatory processes that might take place within these compartments that are less than ∼1 µm in size. Indeed, most of these varicosities are non-synaptic release sites and it remains unclear whether this results in less stringent requirement of the spatial organization of the molecular machinery as compared to e.g. glutamatergic synapses (Choquet and Hosy, 2020; Groc and Choquet, 2020; Liu and Kaeser, 2019). Here, we utilize several super-resolution techniques to provide detailed, quantitative insights into how a key membrane protein, DAT, is subject to nanoscale regulation within these varicosities. This regulation involves Ca^2+^-dependent movements of DAT in and out of PIP2-enriched nanodomains in the plasma membrane, which are governed by NMDA receptor and D2-autoreceptor activity, and likely involves shift of DAT between outward and inward-facing conformations.

To firmly substantiate that DAT is distributed into nanodomains in the plasma membrane of DA neurons, we applied a range of different super-resolution microscopy techniques to both cultured neurons and striatal slices. A varied label density cluster analysis (Baumgart *et al*., 2016) supported that our dSTORM procedure reports true nanodomains, which was further confirmed by ExM, excluding potential artifacts from multiple countings of the same fluorophore. A rough estimate of the number of DAT molecules suggested the presence of 50-100 DAT molecules within each domain, which is in the same range estimated for AMPA receptor domains in the postsynapse (∼25 molecules) (Goncalves et al., 2020). Importantly, by using the novel fluorescent cocaine analogue DG3-63 and analyzing the effect of conformationally biased ligands including noribogaine, JHW007 and nomifensine, we were able to obtain evidence that DAT displays a conformational bias depending on whether it is present in a nanodomain or not. This was further supported by analyzing the disease-associated DAT mutant D421N that displays lowered DA affinity and is biased towards an inward-facing conformation. We should nonetheless note that the distribution of D421N could be affected by the cation leak and anomalous DA efflux that previously have been associated with this mutant (Herborg *et al*., 2018). However, it is appealing to envision that a distribution of DAT between a nanoclustered, inward-facing population of transporter molecules and an unclustered, “free” outward-facing population of DAT molecules represents a means by which DAT activity can be regulated on a very fast time scale. A possible question is whether this dynamic localization of DAT to nanodomains involves changes in DAT oligomerization; however, we find this unlikely considering recent data supporting the existence in the plasma membrane of DA dimers that are highly stable even over the course of minutes (Das et al., 2019). Given the large fraction of DAT present in nanodomains It is also unlikely that DAT in nanodomains just constitutes DAT preparing for internalization. But DAT may internalize from the nanodomains as the inward-facing conformation previously has been associated with DAT internalization (Sorkina et al., 2009), which suggests that at least some internalization takes place from the DAT nanodomains.

Of major importance, we validated in striatal slices the dispersing effect of NMDA on DAT nanodomains by employing a novel Voronoi tessellation-based collection algorithm. In addition to support that our findings in cultured DA neurons can be translated to intact tissue, the algorithm offers possibilities for future detailed, quantitative analyses of protein distribution in neuronal terminals in intact brain tissue. Another important observation was that both acute excitatory input, i.e. activation of NMDA receptors, and sustained alterations of neuronal excitability, as evidenced by expression of either mNaChBac or Kir2.1 (Lin *et al*., 2010; Xue *et al*., 2014) in DA neurons, affect the nanoscale distribution of DAT in presynaptic varicosities. We also observed a marked role of presynaptic D2R activity, further suggesting that the nanoscopic distribution of a membrane protein, such as DAT, is continuously adapted by several mechanisms to the immediate activity level of the neurons in which it is expressed. It may be considered that the effects of the pharmacological manipulations on DAT clustering are quantitatively modest. However, the effects are highly significant, as well as it should be considered that the numerous varicosities included in each quantification may be very heterogenous and possibly differentially sensitive to treatments. Such heterogeneity is directly supported by the finding that only a subset of DA terminals appeared active at any given time (Pereira et al., 2016). Future efforts should further clarify this interesting issue.

Our data revealed a striking co-localization of DAT and PIP2 nanodomains, as well as a dispersing effect of NMDA and haloperidol treatment on both types of domains. We also found that the effect of haloperidol and NMDA on DAT nanodomains not only required opening of VGCCs but also was contingent on the presence Ca^2+^. Interestingly, an increase in cytosolic Ca^2+^ was previously shown to release membrane protein sequestered PIP2, which might indicate a possible PIP2-dependent Ca^2+^-regulation of DAT nanodomain localization (McLaughlin and Murray, 2005). Indeed, the anionic lipid is known to form nanodomains that have been suggested to be important for membrane protein nanoclustering, including STX-1 (van den Bogaart *et al*., 2011). Also, PIP2 is known to regulate many cellular processes including ion channel and transporter function (Hamilton et al., 2014; Suh and Hille, 2008). Both the serotonin transporter (SERT) and DAT bind PIP2 (Buchmayer et al., 2013; Hamilton *et al*., 2014), and for DAT it was suggested that PIP2 binds to the DAT N-terminus and induces a structural change that encourages phosphorylation and a reverse transport mode of DAT critical for amphetamine-induced DA efflux (Belovich *et al*., 2019; Hamilton *et al*., 2014). This is interesting in relation to our data that support an inward-facing configuration of nanodomain localized DAT and thereby presumably a more efflux prone transporter (Robertson et al., 2009). Of note, our previous study suggested a dependency on cholesterol for DAT nanodomain distribution (Rahbek-Clemmensen *et al*., 2017). Furthermore, previous data have indicated that cholesterol binds to DAT and regulates its function by promoting an outward-facing conformation (Hong and Amara, 2010). A conceivable explanation is that cholesterol serves to compartmentalize the membrane to enable a nanoclustered co-distribution of DAT and PIP2 (Carquin et al., 2016; Hwang et al., 1995), and accordingly that cholesterol rather interacts with unclustered, outward-facing DAT. It is even possible that rapid movements of DAT in and out of nanodomains parallels the transport cycle shifting the transporter back and forth from an outward-facing, substrate binding, conformation and unclustered distribution to an inward-facing, substrate releasing (or efflux prone), conformation and clustered distribution. Indeed, our single particle tracking data (Spt-STORM) show that the mean jump distance for a given particle is r∼110 nm in a 16 ms time frame (Figure 5M), suggesting that a single DAT molecule moves 7-14 µm during a single transport cycle of 1-2 sec (Prasad and Amara, 2001). Thus, the movements in the membrane are fast enough for DAT molecules to move in and out of nanodomains during a single transport cycle. Fast movements of DAT have also been supported with classical single particle tracking techniques (Guthrie *et al*., 2020) although use of large quantum dots for labeling suggested somewhat slower diffusion speeds (Thal et al., 2019).

Previous data have indicated that stimulation of D2-autoreceptors of both the D2 and D3 subtype enhance DAT activity (Bolan et al., 2007; Castro-Hernandez et al., 2015; Lee et al., 2007; Meiergerd et al., 1993; Parsons et al., 1993; Zapata et al., 2007). Here we provide evidence that in the DA neuronal cultures the D2-autoreceptor is preoccupied with DA and that this promotes DAT nanoclustering may be consistent with such an enhancement; that is, if D2-autoreceptor activation causes an increase in DAT activity, this would expectably lead to a shift of the DAT conformational equilibrium towards a more inward-facing conformation and, hence, a likely shift towards a higher degree of nanodomain localization. However, our data may challenge that the regulation of DAT by D2-autoreceptors involves a direct interaction as previously suggested (Bolan *et al*., 2007; Castro-Hernandez *et al*., 2015; Lee *et al*., 2007). Although the D2-autoreceptor, like DAT, according to our dSTORM analyses, are distributed into nanodomains, we observed only modest co-localization between the two proteins in the DA terminals. The nanodomains responded, nonetheless, like DAT to NMDA and haloperidol, suggesting that excitatory input promotes nanodomain dispersal, while D2 autoreceptor activation promotes nanoclustering. We speculate that this increased clustering upon activation reflects assembly of the receptor with intracellular signaling complexes and/or dynamics relating to desensitization and/or internalization.

DAT nanodomains showed rather little co-localization with nanodomains of STX-1 and clathrin that importantly also responded differently to NMDA. These differential effects support that the changes we see for DAT, the D2-autoreceptor and PIP2 most likely reflect true molecular redistributions and unlikely are the result of non-specific changes. The decrease in STX-1 nanocluster size and increased localization within dense clusters in response to NMDA is consistent with other studies indicating that breakup of STX-1 nanodomains, correlating with vesicle release, takes place, while at the same time more STX-1 is being brought to the sites of release (Maidorn et al., 2019; Padmanabhan *et al*., 2020). Of interest, the Voronoi tesselated distributions showed that NMDA slightly increased localization of STX1 to unclustered DAT (i.e. the larger tessellated areas). This suggests that the previously described direct interaction between DAT and STX-1, which have been proposed to be important for amphetamine-induced efflux (Binda *et al*., 2008), may take place between the unclustered fractions of the two proteins.

Summarized, by application of single-molecule sensitive super-resolution imaging we show compelling evidence that the molecular machinery supporting volume transmission mediated by a neuromodulator like DA is subject to spatial and temporal nanoscale regulation with putative major impact on synaptic function. The conclusion is substantiated by detailed studies of DAT that according to our results is subject to remarkable nanoscale regulatory processes governed by excitatory input, membrane potential, Ca^2+^ and activity of the presynaptic D2 receptor. Taken together, the present results not only constitute a new framework for further deciphering the nanoscale architecture of presynaptic release sites and the underlying functional implications for the DA system but also add to the growing acknowledgment of nanoscopic regulation as a still poorly explored area of fundamental importance for neuronal signaling.

## Supporting information

Control

NMDA

Haloperidol

## ACKNOWLEDGEMENT

We thank Ralph Götz for support with expansion microscopy and Dr. Emiliana Borrelli for providing tissue from D2R knock-out mice. The work was supported by the Lundbeck Foundation grants R266-2017-4331 (UG), R276-2018-792 (UG), R230-2016-3154 (M.D.L.), R181-2014-3090 (FH), R303-2018-3540 (F.H.) and R231-2016-2481-5 (ATS), Independent Research Fund Denmark – Medical Sciences (U.G. 7016-00325B), and the NIDA-Intramural Research Program Z1A DA000610 (AHN and DAG).

## AUTHOR CONTRIBUTIONS

M.D.L. performed all the experiments and wrote the analyses not hereafter mentioned; A.L.E. devised the cluster size algorithm, identified mRNA expression data and performed D2 receptor histology; A.T.S. performed electrophysiology and generated the construct encoding Cre recombinase under control of a truncated TH promotor; N.O.H. performed the D421N CAD cell experiments; S.H.J. cloned Kir2.1 and mNaChBac plasmids and made non-commercially available viruses; D.A.G. and A.H.N. invented and synthesized the fluorescent cocaine analogs; J.F.S. optimized the DA neuron culture protocol; M.D.L., F.H., and U.G. conceptualized the study, designed the research and interpreted data; U.G., F.H., M.S., and C.W. supervised the research. M.D.L., U.G., and F.H. wrote the paper with contributions from M.S. and all the other authors.

## LEGENDS TO SUPPLEMENTARY FIGURES

**Figure S1.**
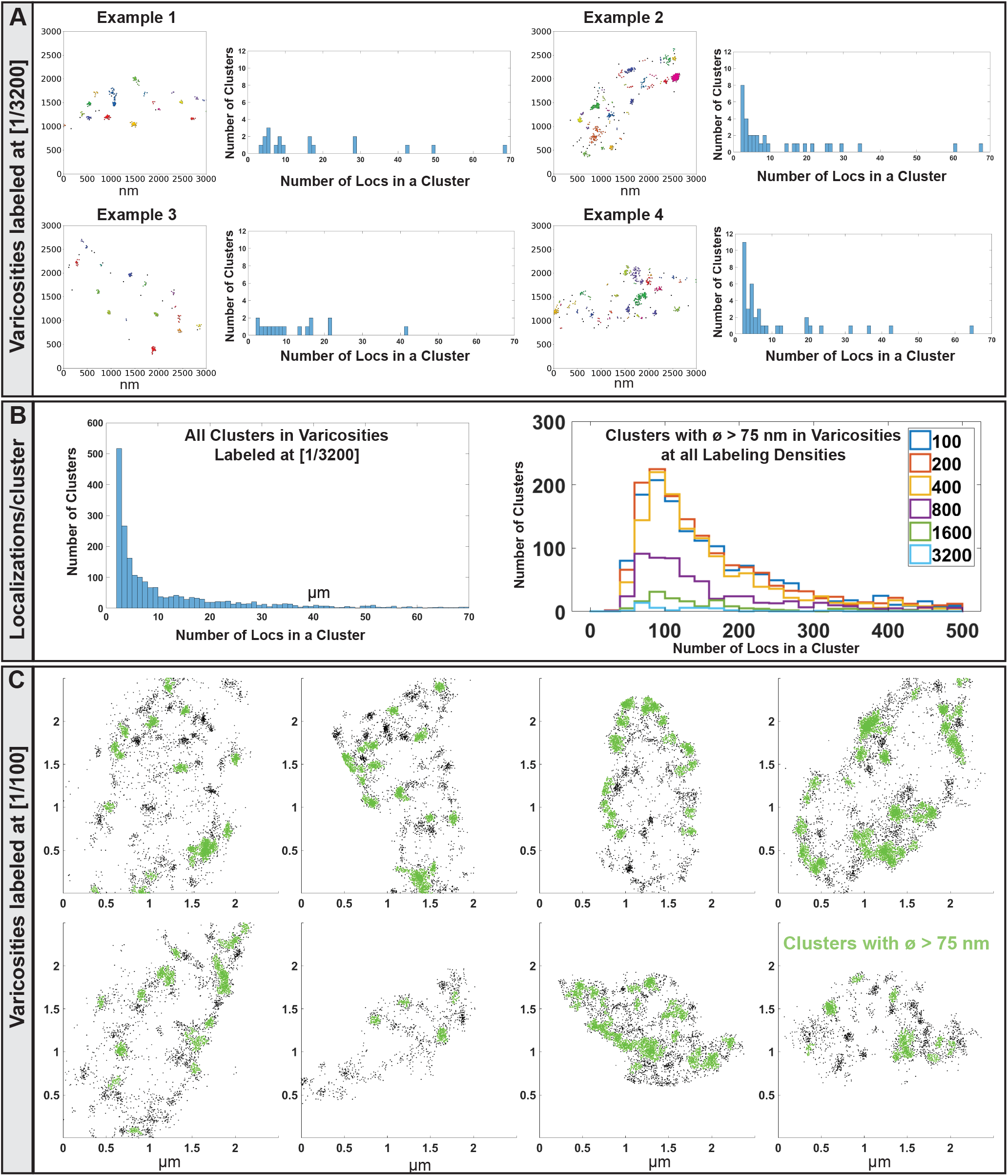
Estimation of protein copy number in DAT clusters (Related to Figure 1.) (A) To estimate how many DAT molecules that are present in an average DAT nanodomain defined by the DBSCAN analysis, we used a protocol modified from (Ehmann *et al*., 2014; Siddig *et al*., 2020) and labeled cultured DA neurons with a twofold dilution row (1:3200-1:100) of primary Mab369 DAT antibody, followed by labeling with Alexa647-conjugated secondary antibody and dSTORM imaging. The figures show four example varicosities labeled with lowest level of primary DAT antibody (1/3200 dilution). Individual nanodomains/clusters are shown by separate colors. For each cluster the number of localizations was counted, as shown by the histogram to right of each example. (B) The clustering analysis was performed on 85 varicosities labeled with 1/3200 dilution of primary antibody and the total number of localizations per cluster for the entire set is shown in the left panel. The graph indicates that the minimum number of localizations in a cluster as a conservative estimate is 3 at this extreme antibody dilution and thus that 3 localizations likely may arise from a single primary antibody in this set of images. We next counted the number of localizations found in varicosities labeled with increasing concentrations of primary antibody (1/100, 1/200, 1/400, 1/800, 1/1600, 1/3200 as indicated by color) and counted the number of localizations present within clusters that has a diameter greater than 75 nm (right panel). The resulting distributions, shown color-coded, peaked at an estimated 100 localizations. It follows that estimated ∼33 copies of the monocolonal antibody would be present that in turn each can label up to two DAT molecules. Accordingly, a rough estimate says that an average cluster contains 50-100 DAT molecules. (C) Example varicosities showing DAT localizations in green in clusters with a diameter larger than 75 nm using a 1/100 dilution of primary DAT antibody.

**Figure S2.**
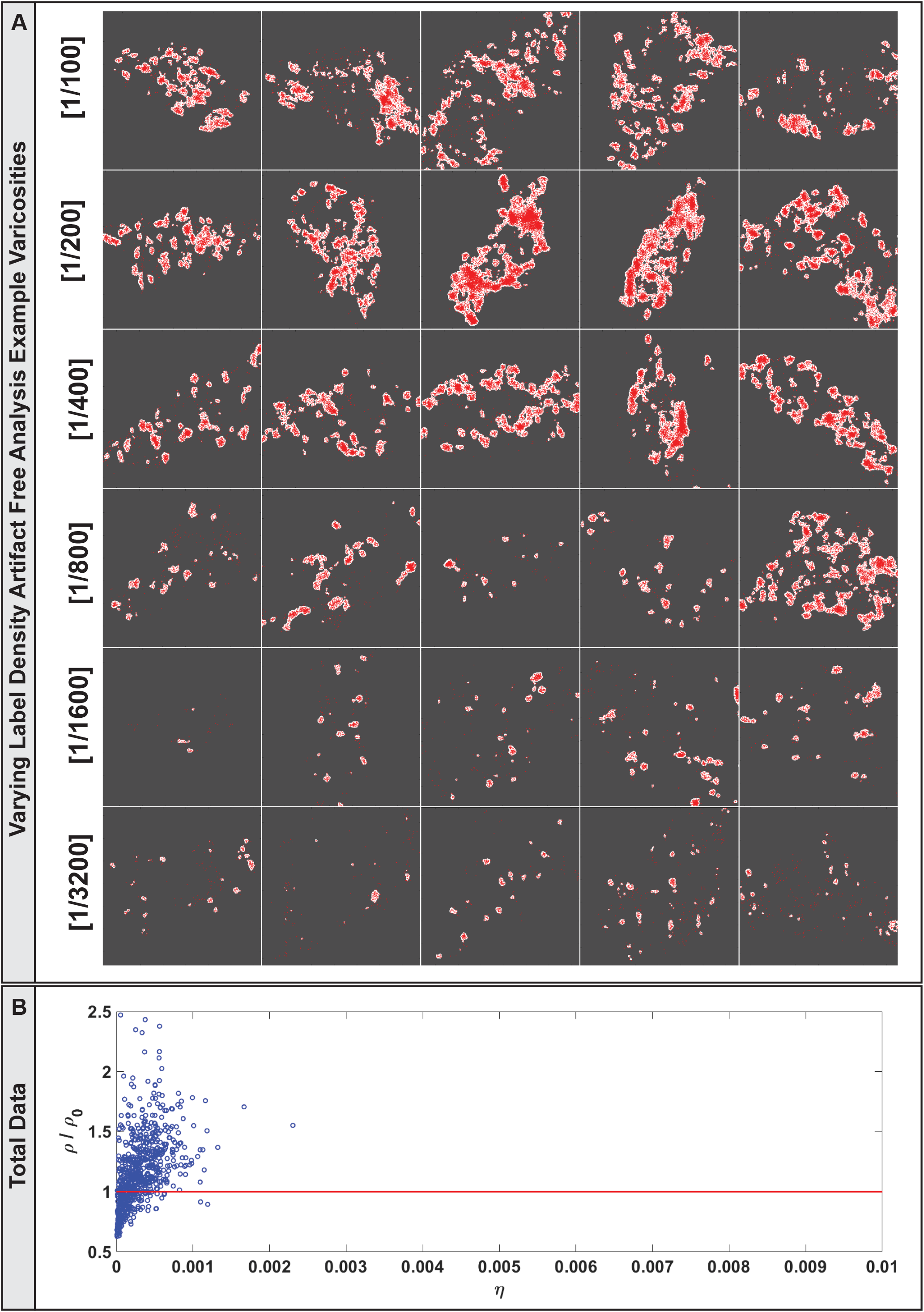
Cluster analysis by varied label density cluster verification (Related to Figure 1.) (A) Example dSTORM images of varicosities from DA neuronal cultures labeled with DAT primary antibody at indicated dilutions. (B) Localization present in clusters (ρ/ρ0) was compared to the relative area of the ROI covered by a cluster mask (η), with each data point being a single varicosity (Baumgart *et al*., 2016). The red line indicates a reference curve from a random distribution and thus the data strongly support a true clustered distribution of DAT localizations into nanodomains and accordingly that the clustering of the DAT signal observed is not the result of multiple observations of single fluorophores in agreement with our previous data in DAT expressing CAD cells (Rahbek-Clemmensen *et al*., 2017).

**Figure S3.**
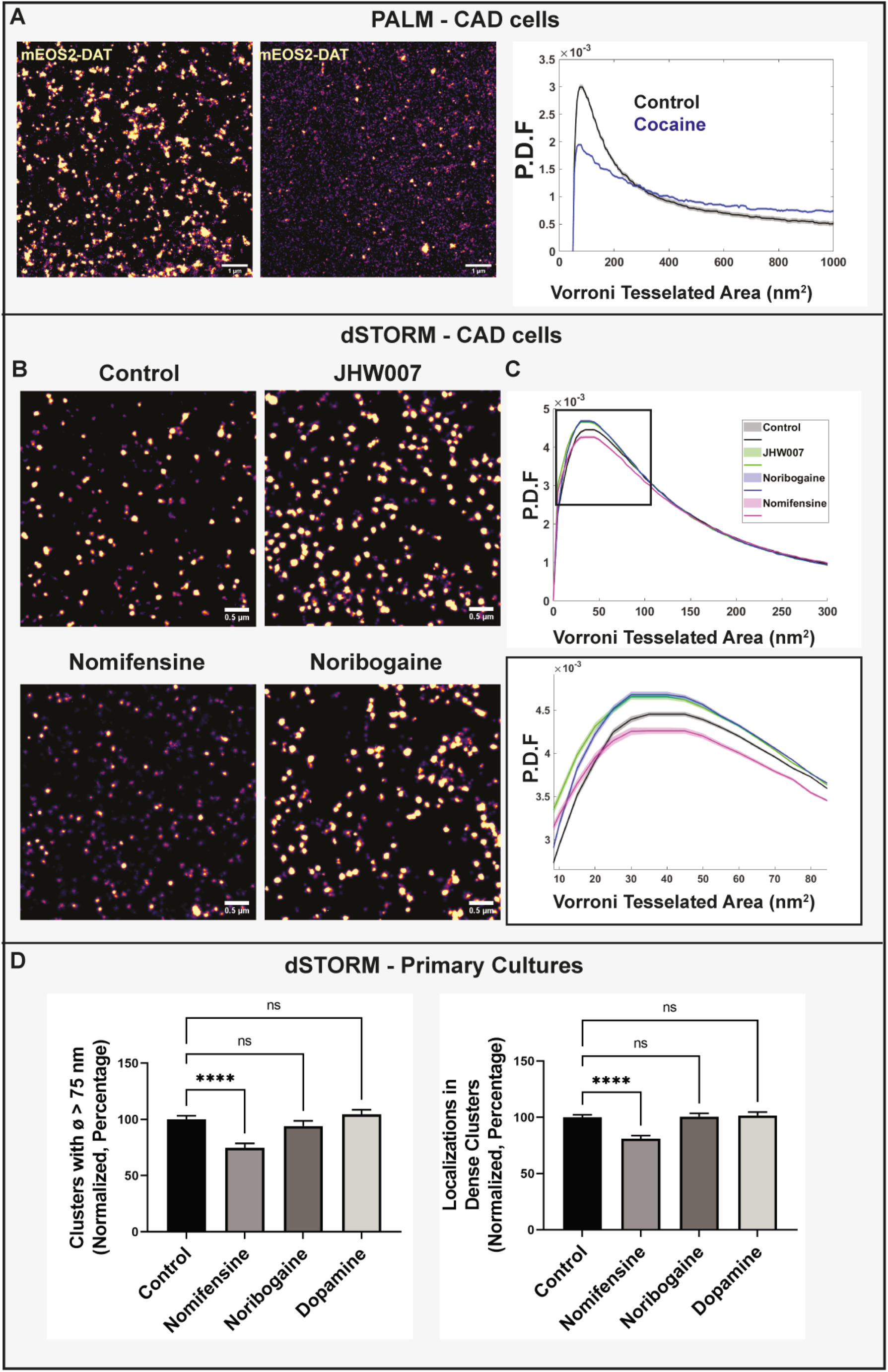
Effects of DAT ligands on DAT clustering in transfected CAD cells and DA neurons (Related to Figure 2). (A) Left, Example PALM/TIRF images of the mEOS2 DAT expressed in plasma membrane of CAD cell with and without incubation with an excess of cocaine. The experiment was done while examining DG3-63’s association with unclustered DAT where unlabeled cocaine was applied to the CAD cells to compete with DG3-63 signal and to show its specificity. The excess cocaine caused a declustering of DAT. Right, Probability density function (P.D.F.) for the Voronoi tessellated areas averaged by varicosity showing a dramatic reduction in the clustering of DAT following cocaine application. Data from 16 control cells (3 transfections) and 17 treated cells (2 transfections). (B) Example dSTORM images visualizing WT DAT expressed in CAD cells after 10 min treatments with JHW007 (20 µM), nomifensine (20 µM), noribogaine (20 µM) or vehicle (control) prior to fixation. (C) Probability density function (P.D.F.) for the Voronoi tessellated areas showing increased clustering of DAT in response noribogaine, JHW007 and decreased clustering in response to nomifensine. Data from 33 control images, 32 noribogaine images, 35 nomifensine images, and 35 JHW007 images. Samples originated from 3 transfections of CAD cells. Curves are means with S.E. indicated by shaded areas. (D) Effect of 5 min treatment of DA primary cultures (prior to fixation and dSTORM imaging) with nomifensine (20 µM), noribogaine (20 µM) or DA (200 nM). Left, Normalized fraction of clusters >75 nm (in %). Right, Normalized fraction of DAT localizations in dense clusters (in %), means ± S.E.; data are from 222 control, 119 nomifensine, 97 noribogaine, and 140 dopamine varicosities obtained in 6 sets of experiments (45 control, 42 nomifensine, 39 noribogaine pictures and 47 DA pictures); one-way ANOVA, ****p<0.0001, n.s., not significant.

**Figure S4.**
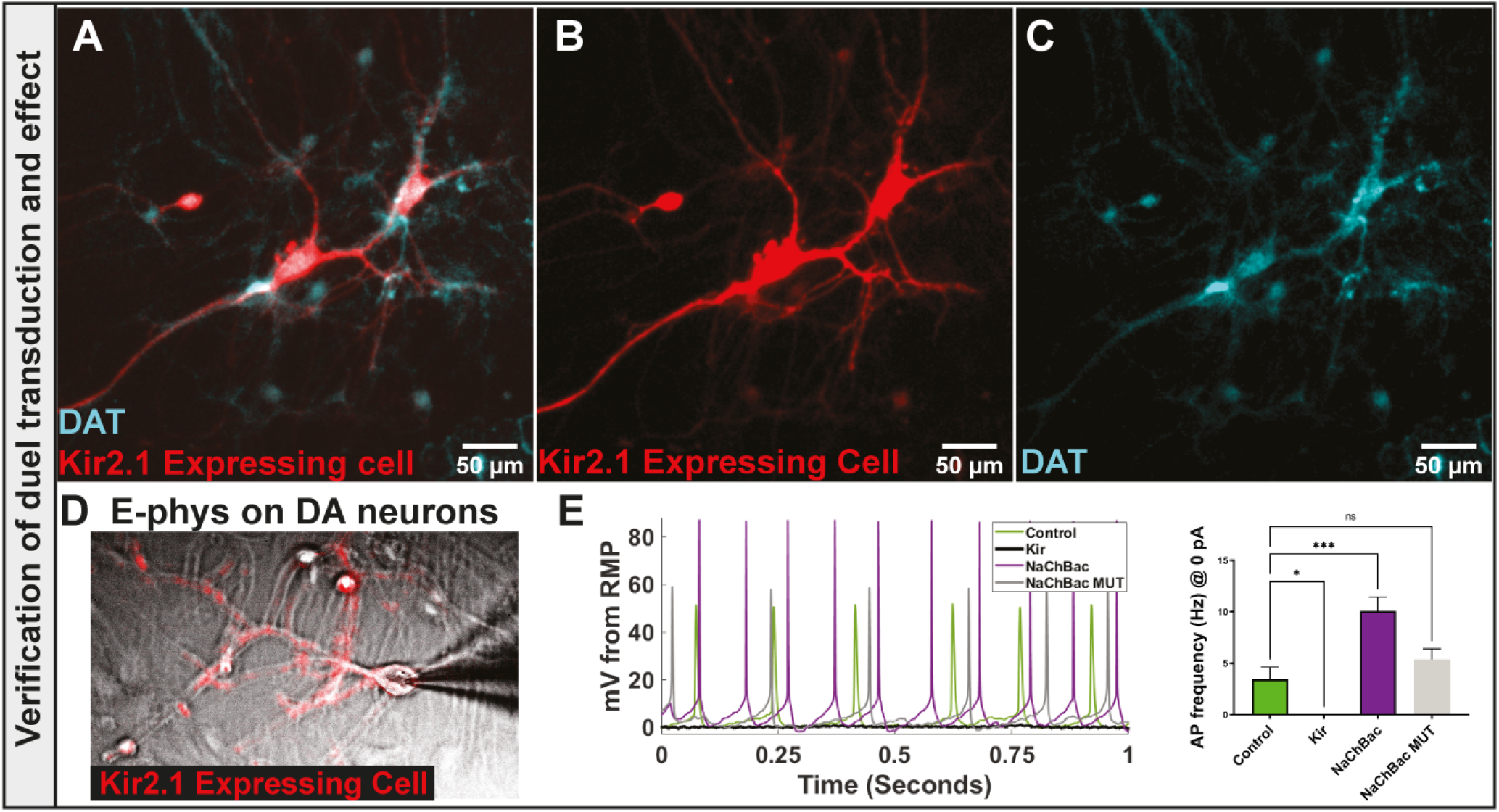
Overexpression in DA neurons of mNaChBac increases while Kir2.1 decreases neuronal excitability (Related to Figure 4). (A-C) Verification of selective expression of bacterial ion channels in DA neurons by a dual viral transduction protocol. The cultured DA neurons were co-infected with an AAV encoding either Cre-dependent mNaChBac or Kir2.1 and with an AAV encoding Cre recombinase under control of a truncated tyrosine hydroxylase (TH) promoter (AAV-pTH-iCre-WPREpA) (see Methods). Images show a putative Kir2.1 expressing DA neuron immunostained for the co-expressed protein mKate2 (red) and for DAT (blue). (A) Widefield image with merged DAT and Kir2.1 signal; B, Kir2.1 signal; C, DAT signal. (D) Whole-cell patch-clamp of a DA neuron expressing Kir2.1. Merged image showing IR-DIC and mKate2 signal of a DA neuron and a patch electrode approaching the cell soma. (E) Left, Current-clamp recording mode (at 0 pA) showing increased firing activity of mNaChBac expressing DA neuron (purple) and essentially no firing activity of Kir2.1 expressing DA neuron (black) as compared to control neuron (green). Right, Action potential frequency (Hz) in control (n=6), mNaChBac (n=4), mNaChBacMUT (n=3) and Kir2.1 (n=5) expressing neurons, means ± S.E.; one-way ANOVA with Dunnett’s multiple comparison test, **p<0.01.

**Figure S5.**
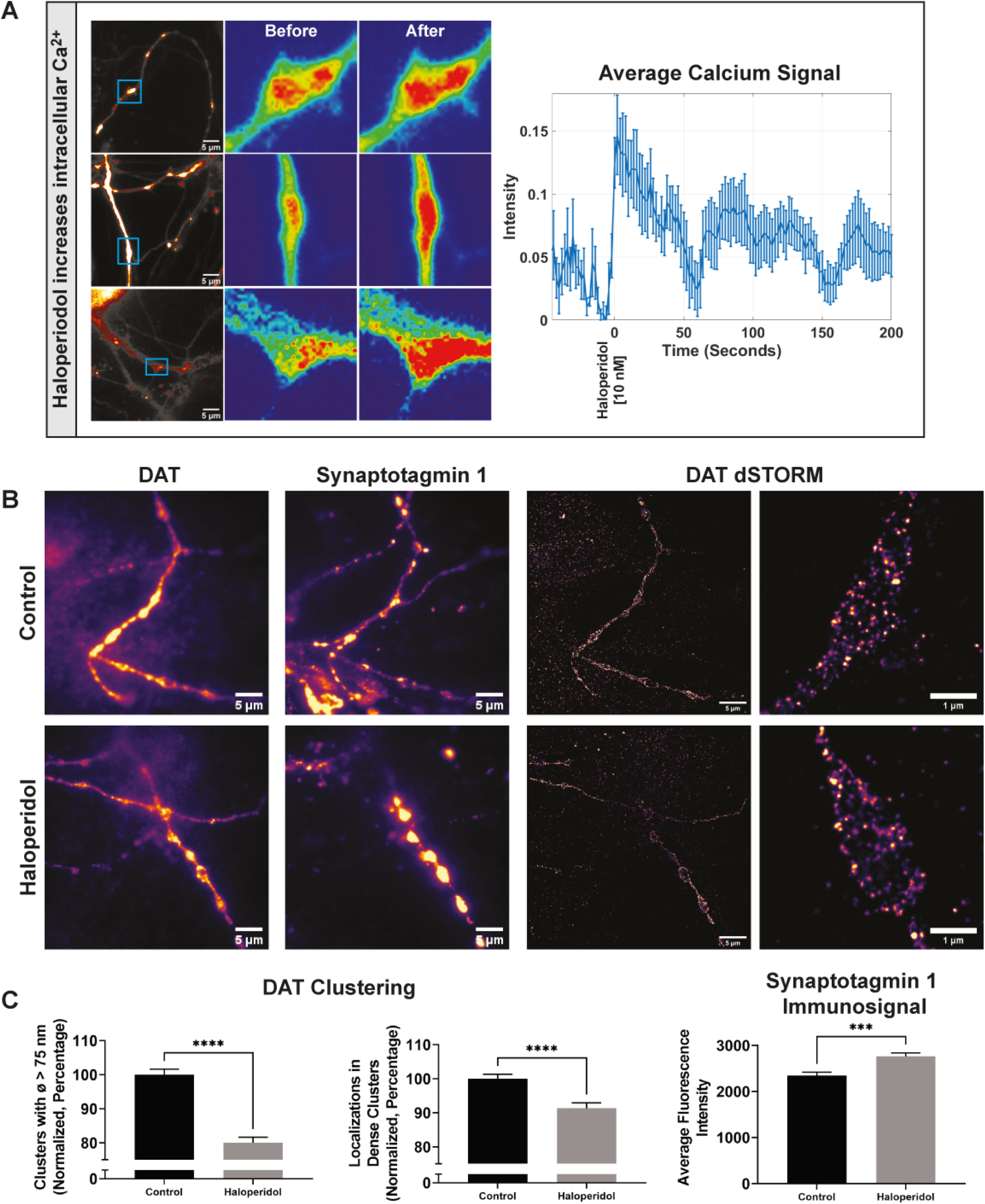
The effect of the D2R antagonist haloperidol on cytosolic Ca^2+^ and synaptotagmin labeling in terminals of cultured DA neurons (Related to Figure 5). (A) Left, Live imaging of terminals from DA neurons expressing the genetically Ca^2+^ sensor jRGECO (three example neurons shown). Right images are enlarged from boxed regions shown in left images. The neurons are, as indicated, shown both immediately before and 200 sec after exposure to 10 nM haloperidol. Right, Averaged fluorescence jRGECO signal over time measured for 6 presynaptic terminals showing how haloperidol (10 nM) cause a rapid rise in cytosolic Ca^2+^. (B) Haloperidol decreases DAT clustering while increasing synaptotagmin 1 labeling in DAT positive varicosities. DA cultures were incubated prior to fixation and immunolabeling for 10 min in the presence or absence of 10 nM haloperidol with antibody targeting the luminal end of synaptotagmin 1. Left, Representative widefield images of DAT and synaptosome with and without haloperidol treatment. Right, Representative DAT dSTORM images with and without haloperidol treatment. Right images are close-ups of the left images. (C) Left, Normalized fraction of DAT localizations in large clusters (>75 nm in diameter) (in %); Middle, normalized fraction of DAT localizations in dense clusters (>80 localizations, radius 50 nm) (in %); Right, synaptostagmin 1 immunofluorescence per varicosity. Data are means ±S.E. and obtained from 429 control varicosities and 464 haloperidol varicosities, across sets of neuronal cultures, one-way ANOVA, ****p<0.0001, ***p<0.001.

**Figure S6.**
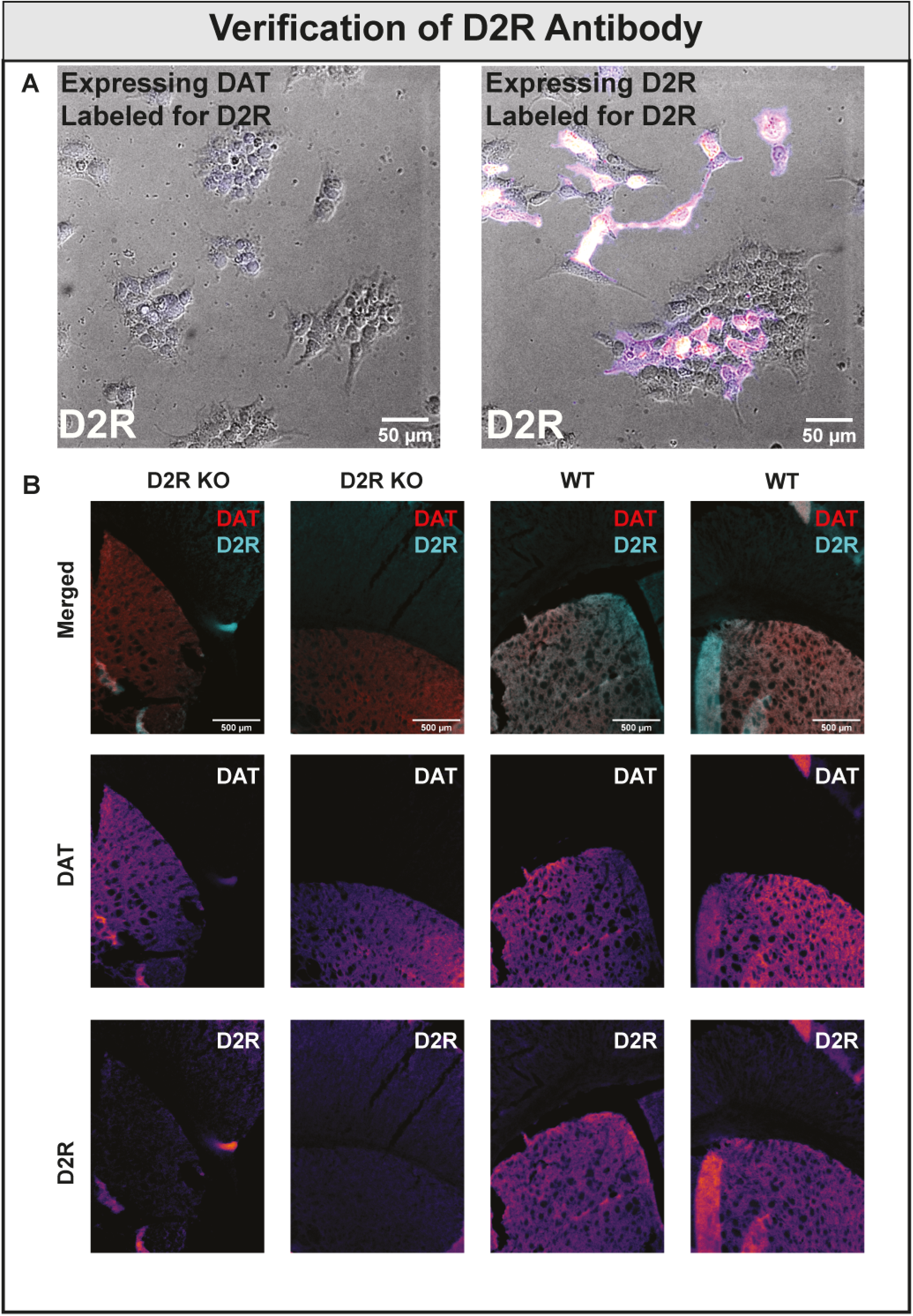
Verification of D2R antibody (Related to Figure 7). (A) Verification of D2R antibody in transfected heterologous cells. HEK293 cell were transiently transfected with the D2R short isoform or DAT. Widefield microscopy show D2R immunosignal in some but not all HEK 293 cells transiently expressing the D2R short isoform. No signal was observed in DAT expressing HEK cells stained with D2R antibody. (B) Verification of D2R antibody in slices from WT and D2R KO mice. Striatal brain slices from D2R KO mice and WT litter mates (Maldonado et al., 1997) were stained for DAT and D2. While a clear immunosignal demarcating striatum was seen for D2 in WT mice, this was absent in the knock-out mice.

**Figure S7.**
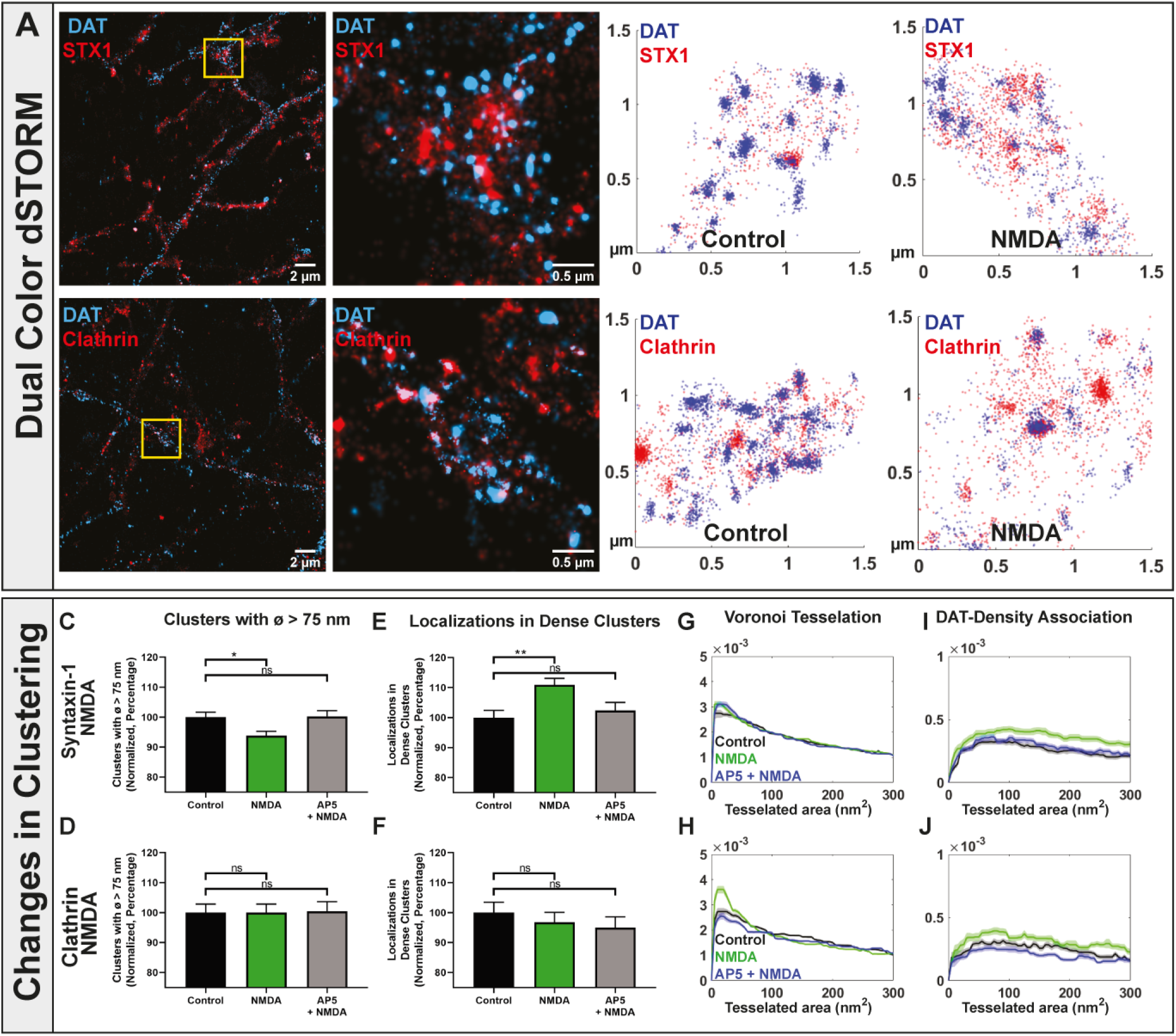
Analysis of STX-1 and clathrin nanodomains (Related to Figures 6 and 7). (A) Example dual-color dSTORM images of syntaxin-1 (STX-1) (red, CF56) and DAT (blue, Alexa647) (upper left panels), and clathrin (red, CF56) and DAT (blue, Alexa647) (lower left panels) in DA neuronal extension (left) with close-up of varicosity (right) corresponding to boxed region in left image. (B) Example dual-color dSTORM images of varicosities from DA neurons showing localizations for syntaxin-1 (STX-1) (red, CF56) and DAT (blue, Alexa647) (upper right panels), and clathrin (red, CF56) and DAT (blue, Alexa647) (lower right panels), Left, Control; Right, neurons exposed to NMDA (20 µM, 5 min). (C-J) Effect of NMDA on STX-1 and clathrin clustering. Cultured DA neurons were stimulated with vehicle (control), NMDA (20 µM) or NMDA (20 µM) + AP5 (100 µM) before DAT and STX-1/clathrin immunolabeling for dSTORM analysis. (C, D) Normalized fraction of STX-1 or clathrin localizations in large clusters (>75 nm in diameter) (in %) and (E, F), normalized fraction of STX-1 or clathrin localizations in dense clusters (>80 localizations, radius 50 nm) (in %), means ±S.E., one-way ANOVA, **p<0.01, *p<0.05, n.s., not significant. (G, H) Probability density functions (P.D.F.) (color-coded as in the bar diagrams) for the Voronoi tessellated areas averaged by varicosity (G, STX-1; H, Clathrin). (I, J) Probability density functions (P.D.F.) (color-coded as in the bar diagrams) of the association of of STX-1 or clathrin with DAT clusters determined by Voronoi tessellation-based association (see Figure 2). Data are from 3 cultures and 217 control, 278 NMDA, 183 NMDA + AP5 varicosities (STX-1); and 98 control, 98 NMDA, 108 NMDA + AP5 varicosities (clathrin).

**Figure S8.**
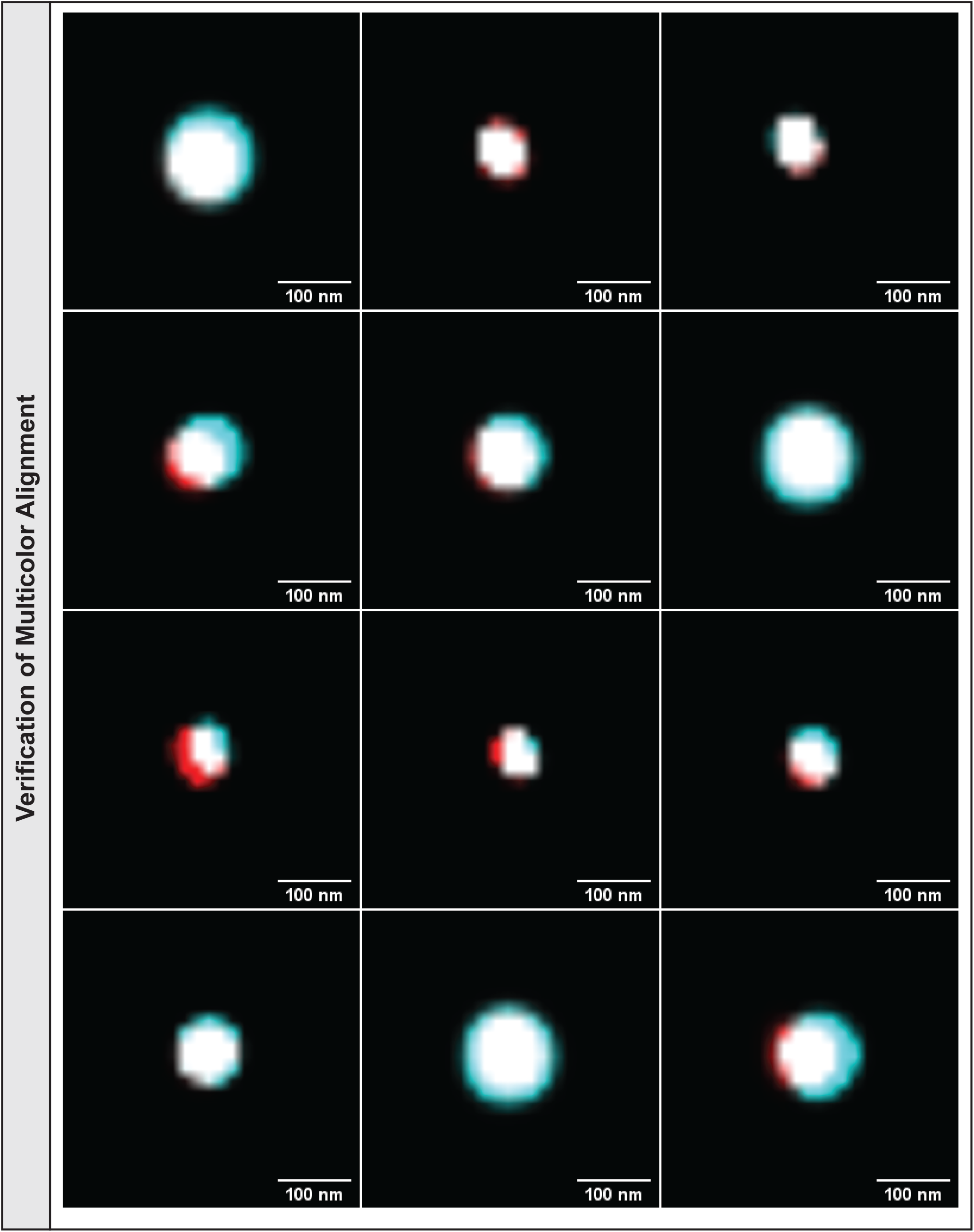
Validation of the multicolor alignment of our microscope (Related to Star Methods). To verify the multicolor alignment of our microscope, dSTORM images of 100 nm Tetraspek beads were taken. Localizations were fit to widefield illuminated Tetraspec beads and processed as our dSTORM data is processed. Localization data was generated into images as such to show how well aligned the multicolor signal is in our system.

## LEGENDS TO SUPPLEMENTARY MOVIES

**Movie S1. 3D-STORM of DA neuronal varicosity immunostained for DAT (control conditions) (Related to Figures 4 and 5).** Color-code is based on density where red localizations are defined as over 300 localizations within 100 nm radius, white as ∼150 localizations within a 100 nm radius, and the remainder in blue.

**Movie S2. 3D-STORM representation of DA neuronal varicosity immunostained for DAT (5 min NMDA treatment) (Related to Figures 4 and 5).** Color-code is based on density where red localizations are defined as over 300 localizations within 100 nm radius, white as ∼150 localizations within a 100 nm radius, and the remainder in blue.

**Movie S3. 3D-STORM representation of DA neuronal varicosity immunostained for DAT (5 min haloperidol treatment) (Related to Figure 5).** Color-code is based on density where red localizations are defined as over 300 localizations within 100 nm radius, white as ∼150 localizations within a 100 nm radius, and the remainder in blue.

## STAR*METHODS

### LEAD CONTACT

Further information and requests for resources and reagents should be directed to and will be fulfilled by the Lead Contact, Ulrik Gether

### MATERIALS AVAILABILITY

All unique reagents generated in this study are available from the Lead Contact. Plasmids generated are available upon request.

### DATA AND CODE AVAILABILITY

Custom written codes used for the data analysis can be found at: https://github.com/GetherLab/Super-Resolution-Data-Analysis. The datasets supporting the current study have not been deposited in a public repository because of the file size but are available from the corresponding author on request.

### EXPERIMENTAL MODEL AND SUBJECT DETAILS

#### Primary cultures of DA neurons with cortical glia cells

Cultured glial cells were obtained from cortex and cultured DA neurons were obtained from the midbrain of P1-P3 Wistar rats (Charles River, Germany). Gender was mixed as the cultures were generated from a mixed sex populations of rat pups. For preparation of cultured glial cells, brains were removed from euthanized P1-P3 pups and placed in phosphate-buffered saline (PBS) on ice. The cortex of each brain was isolated and cut into millimeter sized sections and collected in cold dissection medium (Hanks’ Balanced Salt Solution with added Sodium Pyruvate (1 mM), Penicillin-Streptomycin (Sigma P0781, 200 U/L), 10 mM HEPES, and 0.54% glucose)). After collecting all, the dissection medium was replaced with glia cell medium (DMEM with HEPES “1965”, FBS (10%), Penicillin-Streptomycin (60 U/L)). The tissue was mechanically titrated through pipetting up and down using a flame-polished Pasteur pipette followed by straining through a 70 µm cell trainer and centrifugation for 10 min at 200-300 x g. The pellet was resuspended in 10 mL glia medium and strained again before a second centrifugation for 10 min at 200-300 g and removal of the supernatant. The pellet was resuspended and seeded into large T175 flasks at approximately 1.5 brains per flask with added glial medium to a total of 20 mL. Glia cells were grown in an incubator at 37° C with 5% CO_2_ until they were 70% confluent. The cells were either frozen in heat-inactivated serum containing 10% DMSO for later use or applied to ethanol-cleaned and poly-D-lysine-coated coverslips and incubated for 1 week prior to DA neuron addition. On the day before DA neuron addition, the glial cells were changed to DA neuron medium (Neurobasal A with heat-inactivated FBS (1/100), GlutaMAX (1/100) B-27 (1/50), ascorbic acid (200 µM), Penicillin-Streptomycin (60 U/L), kynurenic acid (0.5 mM)). Midbrain DA neurons were generated using a protocol modified from (Rayport *et al*., 1992). Brains from rat pups (P1-P3) were isolated and placed in PBS on ice. Midbrain sections were retrieved from each brain as the brains were cut while in dissection buffer. The midbrain sections were transferred to papain solution (cysteine (1 mM), DNase (0.1 mg/mL), papain (20 units/mL), NaCl (116 mM), KCl (5.4 mM), CaCl_2_ (1.9 mM), NaHCO3 (26 mM), NaH_2_PO_4_H_2_O (2 mM), MgSO_4_ (1 mM), EDTA (0.5 mM), Glucose (25 mM), kynurenic acid (0.5 mM) in water) for 30 min at 37°C. The solution was replaced with prewarmed DA medium and samples were mechanically titrated with flame polished Pasteur pipettes and the resulting suspension centrifuged at 200-300 x g for 10 min. The pellet was resuspended in DA neuron medium and applied to each culture. Glia-derived neurotrophic factor (GDNF) (100ng/mL of culture solution volume) was applied immediately after seeding of the neurons. Cultures were incubated for 2 to 3 weeks prior to use. All procedures were performed according to institutional guidelines at the Faculty of Health Sciences, University of Copenhagen.

#### Mouse brain slices

Mouse brain slices were obtained from male C57BL/6 mice 12-16 weeks old. The slices were prepared as described in Methods Details. For validation of D2 receptor antibody, slices were obtained from D2 receptor knock-out mice and WT littermates (Maldonado *et al*., 1997). All procedures were performed according to institutional guidelines at the Faculty of Health and Medical Sciences, University of Copenhagen.

#### Heterologous cell culture

Cath.a-differentiated (CAD) cells (Qi et al., 1997) (ATCC CRL-11179 or Sigma-Aldrich 08100805), originating from mouse (B6/D2 F1 hybrid) catecholaminergic neuronal tumor. The cells were maintained in a 1:1 mixture of DMEM 1965 and Ham’s F-12 medium (Invitrogen), both supplemented with 10% FBS and 0.01 mg/mL gentamicin. HEK293 cells (human embryonic kidney cells, ATCC CRL-1573) were grown as described (Hansen *et al*., 2014).

### METHOD DETAILS

#### Molecular biology, virus generation and virus transduction

The cDNA encoding human DAT (synthetic gene kindly provided by Dr. Jonathan Javitch, Columbia University, NY, USA (Loland et al., 2004) was inserted into pmEOS2-C1 (RRID: Addgene#54510) to generate pmEOS2-hDAT C1 encoding hDAT with mEOS2 fused to the N-terminus. Plasmids encoding WT hDAT and D421N hDAT (pRC/CMV hDAT and pRC/CMV hDAT D421N) were generated as described (Herborg *et al*., 2018)). The cDNA encoding the short transcriptional variant of human DA D2 receptor with an N-terminal FLAG tag (SF-D2R-S) was inserted into mammalian expression vector pcDNA 3.1(+) (Invitrogen) (Klewe et al., 2008). The Cre-dependent construct, pAAV-CAG-Flex-NES-jRGECO1a-WPREpA, was obtained from Addgene (RRID: Addgene #100854). The plasmids, pAAV-hSyn-DIO-mNaChBac-T2A-mKate2-WPREpA, pAAV-hSyn-DIO-mNaChBacMUT-T2A-mKate2-WPREpA and pAAV-hSyn-DIO-Kir2.1-T2A-mKate2-WPREpA (Lin *et al*., 2010; Xue *et al*., 2014), were SLIC cloned (Jeong et al., 2012) from pAAV-EF1α-F-FLEX-mNaChBac-T2A-tdTomato (RRID: Addgene #60658) and pAAV-EF1α-F-FLEX-Kir2.1-T2A-tdTomato (RRID: Addgene #60661), respectively, into a Cre-dependent AAV backbone plasmid with a mKate2 fluorophore instead of tdTomato. This was done to make them sufficiently short so that packaging into an AAV capsid would be feasible (ITR to ITR length<4700 kbp). The pAAV-pTH-iCre-WPREpA construct was created and cloned in-house and contains a truncated 488bp rat TH promoter encoding Cre recombinase. In-house generated AAVs were produced using a FuGene6 (Promega) mediated triple plasmid co-transfection method in HEK293t cells. Three days after transfection, cells were harvested and virus purified using an adapted Iodixanol gradient purification protocol (Matsui et al., 2012). Genomic AAV titer was determined by a PicoGreen-based method as described elsewhere (Piedra et al., 2015). AAV2-retro-pTH-iCre mediated expression of Cre-dependent jRGECO1 was verified by fluorescence microscopy and no differences were observed across the different titers used. For transduction, the appropriate virus was applied to primary culture DA neurons at an approximate concentration of 2*10E9 vg/mL at 1 to 3 days after preparation of the neuronal cultures.

#### Transfection of DA neurons

Primary cultures of DA neurons were transfected using Magnetotransfection at ∼14 DIV (Underhill et al., 2014). NeuroMag reagent (OZ biosciences) (8 µL) was mixed with plasmid (4 µg) and DA neuron medium (200 µL). The mixture was incubated for 20 min at RT before it was added to cultures grown in 6-well plates. The culture was positioned on top of a magnetic plate for 15 min after which the culture before it was moved back to the cell incubator and used for immunostaining 24-48 hours later.

#### Immunostaining of primary cultures of DA neurons for dSTORM and PALM

The cell samples were fixed in paraformaldehyde (3%) and washed three times in glycine (20 mM) and NH_4_Cl (50 mM) in PBS. Subsequently, the cells were washed in blocking buffer (5% Donkey serum, 1% BSA in PBS), and incubated in blocking permeabilization buffer (blocking buffer with saponin (0.2%)). Primary antibody was applied in blocking buffer for 60 min, followed by 3–5 min incubation in blocking buffer. Secondary antibody was applied in blocking buffer for 45 min, and the sample was incubated 2x for 5 min in blocking buffer. Samples were washed in PBS twice and post-fixated in paraformaldehyde (3%) for 15 min. Samples were washed twice in glycine (20 mM) and NH_4_Cl (50 mM) in PBS and stored in PBS at 4°C until imaging.

The synaptotagmin 1 feeding experiment was performed as defined by (Truckenbrodt *et al*., 2018). Live DA primary cultures were exposed to synaptotagmin 1 antibody (SS 105 311, Synaptic Systems) at a 1:120 dilution from a 1 mg/mL solution together with haloperidol (10 nM) or vehicle 37°C for one hour. The cultures were carefully washed with 37°C aCSF while avoiding exposure to air before fixation and immunolabeling per the primary culture staining protocol.

#### Preparation and immunostaining of mouse brain slices for dSTORM

Mice were anesthetized with isoflurane and perfused first with PBS followed by 4% paraformaldehyde (PFA) injected directly into the heart. Brains were then placed in 4% PFA for 24 hours, followed by sucrose (30%) for 24 to 48 hours. Subsequently, brains were snap-frozen on powdered dry ice and stored at - 80°C. The brains were sliced using a cryostat (Leica CM3050 S) to obtain 10 μm striatal coronal sections and each slice was immediately fused to a 3-aminopropyltriethoxysilane coated coverslip and let dry at room temperature before being stored in antifreeze (50% glycerol 50% PBS) at −20 °C. For immunostaining, brain sections were first rinsed 3x in PBS and subsequently washed 2x with 15 min incubation in glycine (20 mM) and NH_4_Cl (50 mM) in PBS. This was followed by 30 min wash in 10 mM trisodium citrate, pH 6.0, preheated to 80° C, 4x wash in PBS and then blocking/permeabilization for 30 min in PBS with 5% donkey serum, 1% BSA and 0.3 % Triton X-100. Primary antibody was applied overnight in blocking/permeabilization buffer followed by washing in 5/5/90 min increments and application of secondary antibody overnight in blocking/permeabilization buffer. Sections were again washed in 5/5/90 minute increments in blocking/permeabilization buffer and then with 5/5 min increments with PBS. The sections were post-fixated in paraformaldehyde (3%) and washed twice for 15 min in glycine (20 mM) and NH_4_Cl (50 mM) in PBS. Samples were lastly washed twice in PBS and stored in PBS at 4 °C until imaging.

#### Antibodies and labeling

Secondary antibodies (donkey anti-rat, Jackson 712-005-153, RRID:AB_2340631; donkey anti-mouse, Jackson 715-005-151, RRID:AB_2340759); or donkey anti-rabbit, Jackson 711-005-152, RRID:AB_2340585) were labeled with either NHS-ester conjugated Alexa647 or CF™568 (Biotium, Freemont, CA) and then isolated through Zeba™ Spin Desalting Columns, 40K MWCO (ThermoFisher 87767). This was completed by first washing the column 3x with 300 µL of 100 mM NaHCO_3_ in PBS. Following the washes, 50 µg of antibody in 100 µL of PBS was added and spun at 1500 x g for 2 mins. The flow-through was collected and combined with a 5-fold molar excess of NHS ester fluorophore. This solution was incubated in the dark at RT, shaking, for 2.5 hours. A new spin column was then washed 3 times with NaN--3 (0.02%) in PBS. The antibody sample with the dye was then added to the column and centrifuged for 2 min at 1500 x g. The protein concentration and label concentration of the final product were measured on a NanoDrop 2000 Spectrophotometer (ThermoFisher). Labeled antibodies had between 1 and 2 fluorophores attached.

We used the following antibodies and fluorophores for our imaging analysis:

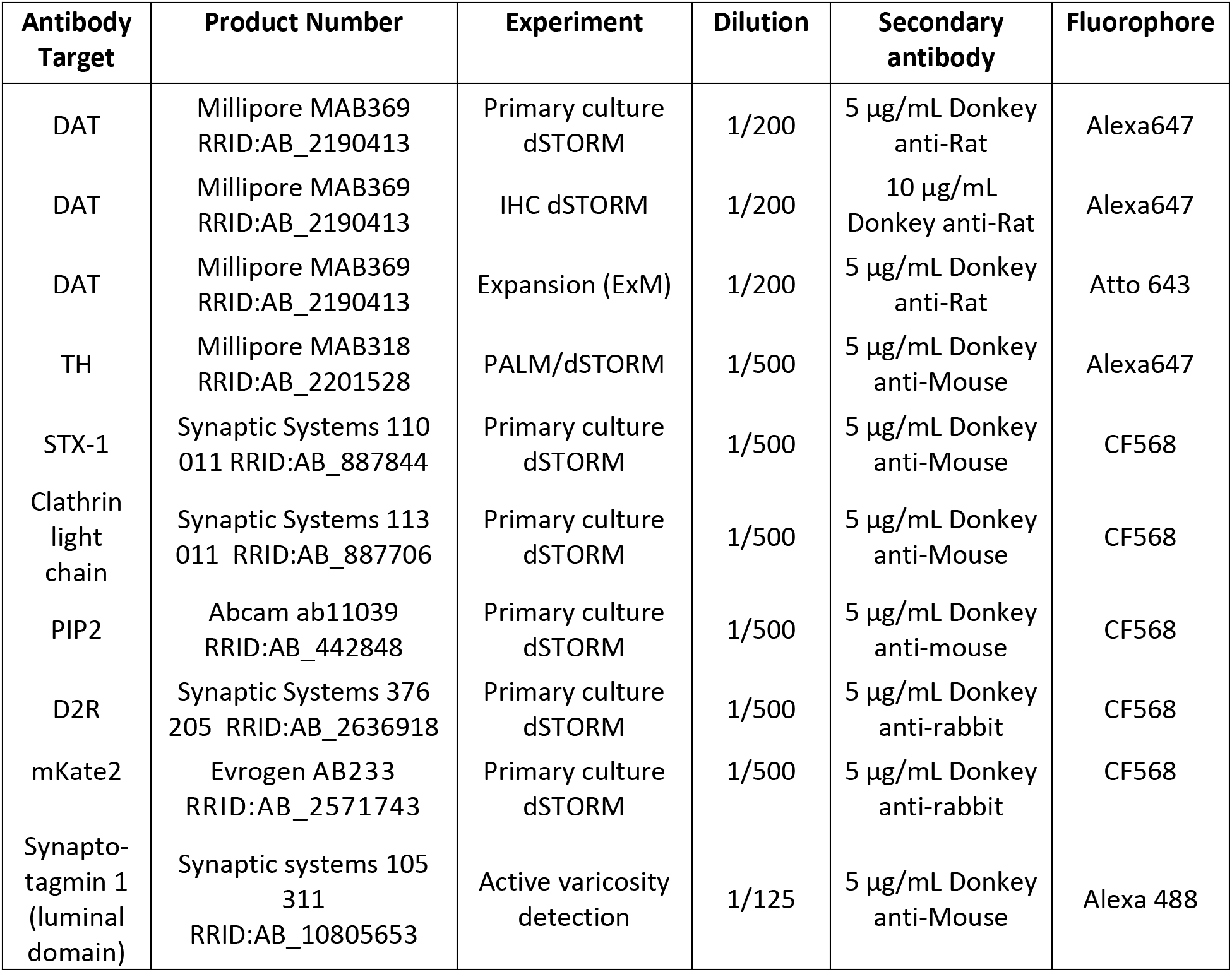

| Antibody Target | Product Number | Experiment | Dilution | Secondary antibody | Fluorophore |
| --- | --- | --- | --- | --- | --- |
| DAT | Millipore MAB369<br>RRID:AB_2190413 | Primary culture<br>dSTORM | 1/200 | 5 µg/mL Donkey<br>anti-Rat | Alexa647 |
| DAT | Millipore MAB369<br>RRID:AB_2190413 | IHC dSTORM | 1/200 | 10 µg/mL<br>Donkey anti-Rat | Alexa647 |
| DAT | Millipore MAB369<br>RRID:AB_2190413 | Expansion (ExM) | 1/200 | 5 µg/mL Donkey<br>anti-Rat | Atto 643 |
| TH | Millipore MAB318<br>RRID:AB_2201528 | PALM/dSTORM | 1/500 | 5 µg/mL Donkey<br>anti-Mouse | Alexa647 |
| STX-1 | Synaptic Systems 110<br>011 RRID:AB_887844 | Primary culture<br>dSTORM | 1/500 | 5 µg/mL Donkey<br>anti-Mouse | CF568 |
| Clathrin<br>light<br>chain | Synaptic Systems 113<br>011 RRID:AB_887706 | Primary culture<br>dSTORM | 1/500 | 5 µg/mL Donkey<br>anti-Mouse | CF568 |
| PIP2 | Abcam ab11039<br>RRID:AB_442848 | Primary culture<br>dSTORM | 1/500 | 5 µg/mL Donkey<br>anti-mouse | CF568 |
| D2R | Synaptic Systems 376<br>205 RRID:AB_2636918 | Primary culture<br>dSTORM | 1/500 | 5 µg/mL Donkey<br>anti-rabbit | CF568 |
| mKate2 | Evrogen AB233<br>RRID:AB_2571743 | Primary culture<br>dSTORM | 1/500 | 5 µg/mL Donkey<br>anti-rabbit | CF568 |
| Synapto-<br>tagmin 1<br>(luminal<br>domain) | Synaptic systems 105<br>311<br>RRID:AB_10805653 | Active varicosity<br>detection | 1/125 | 5 µg/mL Donkey<br>anti-Mouse | Alexa 488 |

#### Expansion Microscopy (ExM)

For ExM, perfused brains were acquired by injecting first PBS and then 4% PFA into the heart of anesthetized mice. Brains were postfixated in 4% PFA for 24 hours and immersed in sucrose solution (30%) for 24 to 48 hours. Coronal brain slices (40 µm) were obtained with a cryostat. The brain slices were stored, free floating, in antifreeze (50% glycerol 50% PBS) at −20°C until use. Brain slices were washed with PBS three times, and then permeabilized in blocking buffer (5% donkey serum, 1% bovine serum albumin, 0.1% Triton X-100, in PBS). Brain slices were incubated overnight with primary antibody in blocking buffer (5% donkey serum, 1% bovine serum albumin, 0.1% Triton X-100, in PBS, pH 7.4). Slices were washed 4x in blocking buffer with 30 min increments and incubated overnight with secondary antibody in blocking buffer at 4°C. Slices were again washed 4x in blocking buffer with 30 min increments. Anchoring treatment was performed with Acryloyl-X-SE (0.1 mg/mL) in PBS at room temperature overnight (Truckenbrodt *et al*., 2019). Gel solution was made by mixing N,N-dimethylacrylamide (1.335 g) and sodium acrylate (0.32g) in 2.85 g of water. Oxygen was purged with nitrogen gas for 40 min. 2.7 mL of the gelling solution were mixed with 0.3 mL 0.036 g/mL solution of potassium persulfate (K_2_S_2_O_8_) solution and purged of oxygen with nitrogen gas for 15 min on ice. Brain slices were transferred to the extra gelling solution and placed flat on poly-D-Lysine coated glass slides. Excess gelling solution was removed, and glass coverslips were placed on the edges of each slice to limit gel dispersion. All work from this point was done in a cold room (4°C). Aliquots of 500 µL of polymerization solution were combined with 2 µL of TEMED was added and the solution vortexed before applying 80 µL to brain slices. A layer of parafilm covered coverslip was placed overtop of the gel to fully surround the gel. The sample was kept in the cold room for 15 min before being brought back out to room temperature. The container with the gels was filled with nitrogen and wet paper to keep humidity and made airtight. Following gelation, samples were carefully cut out from the chamber in a sim-card shape to ensure orientation. Gels were measured in size, and placed in digestion buffer overnight at room temperature (50mM TRIS, 800 mM guanidine HCl, 2 mM CaCl2, 0.5% Triton X-100, proteinase K 8U/mL, pH 8.0). The gels were placed into individual dishes and dialyzed with water many times over a day until they no longer expanded. Imaging was performed the next day on a spinning disk confocal microscope (FEI CorrSight utilizing a C-Apochromat 63x/1.20 W Corr M27 objective). Samples were places dry on poly-D-Lysine coated chambers and water was added to the top of the gel just enough so the gel was hydrated but not so it would move around.

#### dSTORM

For dSTORM we used a buffer containing β-mercaptoehtanol and an enzymatic oxygen scavenger system (10% (w/V) glucose, 1% (V/V) beta-mercaptoethanol, 2 mM cyclooctatetraene, 50 mM Tris-HCl (pH 8), 10 mM NaCl, 34 μg mL^-1^ catalase, 28 μg mL^-1^ glucose oxidase). The imaging was performed with an ECLIPSE Ti-E epifluoresence/TIRF microscope (NIKON, Japan) equipped with 405 nm, 488 nm 561, and 647 nm lasers (Coherent, California, USA). All lasers are individually shuttered and collected in a single fiber to the sample through a 1.49 NA, 100x, apochromat TIRF oil objective (NIKON). Single color imaging was done with a dichroic mirror (z405/488/561/647 rpc) and the emitted light was filtered by a 710/80 nm bandpass filter. For dual-color dSTORM, we used a dichroic mirror with the range 350–412, 485–490, 558–564, and 637–660 nm (97,335 QUAD C-NSTORM C156921). The excitation light was filtered at the wavelengths: 401 ± 24 nm, 488 ± 15 nm, 561 ±15 nm, 647 ± 24 nm. The emitted light was filtered at the wavelengths: 425-475, 505-545, 578-625, and 664-787 nm, and secondly by an extra filter to decrease noise (561 nm Longpass, Edge Basic, F76-561, AHF). A motorized piezo stage controlled by a near-infrared light-adjusted perfect focus system (NIKON) is applied to the system to reduce any sample drift over time in the z-direction. Single-color dSTORM images were constructed from 30,000 frames taken at a 16 ms frame rate. Dual-color dSTORM images were constructed from 20,000 frames for each color taken at a 16 ms frame rate with each color alternating by frame. Photons were collected with an iXon3 897 EM-CCD camera camera (Andor, United Kingdom). Laser powers used were 2.3 kW cm −2 for 647 nm, 1.0 kW cm−2 for 488 nm and for 561 nm. The 405 nm laser was used to incrementally increase blinking behavior at power <0.1 kW cm−2. For 3D-dSTORM, a cylindrical lens was placed before the camera to impart astigmatism.

#### PALM

For PALM imaging of mEos-DAT, we obtained 5,000 consecutive frames with a frame rate of 33 Hz to construct one image. The 488 nm excitation laser was held constant, 0.4 kW cm^-2^, during the capture of the image while the 405 nm activation laser was gradually increased to <0.1 kW cm^-2^.

#### Localization fitting of dSTORM data

2D localizations were fit with ThunderSTORM using local maximum detection with a threshold of 1.5 *std(Wave.F1) and estimator of an integrated gaussian PSF. Sigma was 1.6, fit radius was 3 pixels, and weighted least squares was the method to fit the estimated gaussian to the PSF (Ovesny et al., 2014). 3D localization fitting was performed with fit3Dcspline (Li *et al*., 2018). 3D calibrations were made on 100 nm TetraSpeck beads attached to a coverslip by first placing 50 µL of MgCl2 (1M) on the coverslip and then adding 500 L of water with 0.9 µL of TetraSpeck beads to the coverslip. Drift for all data was corrected with redundant cross correlation through Matlab (Wang et al., 2014). Localizations were filtered for uncertainty being less than 25 nm and then merged for localizations that were detected within 15 nm and 3 frames.

#### Data processing pipeline - dSTORM

To perform necessary localization fitting on the 1349 dSTORM images included in this study, an in-house data processing pipeline was developed to streamline operations. Localizations were fit to the raw data with the aforementioned settings in ThunderSTORM through a self-written ImageJ macro. Although ThunderSTORM can perform the remaining processing steps, a custom python script (utilizing Matlab functions) was written to perform cross correlation, merging and filtering so that redundant cross correlation could be employed. The images were then blinded by labeling them with a random integer and saving the cypher elsewhere. Through a separate custom Matlab script, varicosities were selected from each image though a freehand selection. Varicosities were identified by the local swelling of their structure compared to the rest of the extension. We next used DBSCAN (density based spatial clustering of applications with noise) (Ester *et al*., 1996; Rahbek-Clemmensen *et al*., 2017) to identify clusters by size or for determining localizations in dense clusters through the use of a python script employing the DBSCAN method from the sklearn python library. Voronoi tessellation (Levet *et al*., 2019) was employed through Matlab. The Voronoi based density association algorithm was also written in house and using Matlab.

#### Varicosity identification in dSTORM DA neuron culture data

Automatic varicosity identification could not discern between extension and varicosity, so these were selected through a freehand ROI taken of blinded files. Localization data for all images in a given dataset were blinded by being randomly assigned a number that was stored on a cypher. ROIs were then gathered without the knowledge of the image identity, with this information restored in the analysis of the ROIs.

#### Varicosity identification in brain slice dSTORM data

As this was a computationally intensive process, each image was divided into 100 quadrants in a 10×10 grid. Within each quadrant, the localizations were sub-sampled down to 1000 random localizations, and the Voronoi tessalation was obtained for that section. Tesselated areas were filtered for having an area less than 15000 nm^2^. Tesselated shapes that were touching one another were merged together. Shapes were now filtered for possessing an area greater than 300000 nm^2^. These ROIS were buffered by 50 nm along each edge to smooth the ROI.

#### Clustering analysis of dSTORM data

Cluster size was identified by performing DBSCAN on each varicosity for a radius of 15 nm and 5 localizations being the cluster qualifier. The convex hull was gathered for each cluster to find its area, and the fraction of clusters with a diameter >75 nm was used for the cluster size analysis. Cluster size distributions were not shown as clusters reaching the detection limit of this method are not accurate and cause a false perception on the cluster size distribution. Localizations in dense clusters were found by applying DBSCAN to each varicosity with a radius of 50 nm and 80 localizations being the qualifier. Voronoi tessellation was applied to each varicosity and areas were filtered for sizes between 5 nm2 and 1000 nm2. Probability density functions were created for these distributions.

#### Density Association

Density association was found by performing Voronoi tessellation on target A, and finding all points in target B that were 25 nm from any point in target A. Each of these target B points that possessed A association were mapped to the Voronoi tessellated area from target A they resided in. The probability density functions for target A and the areas from target A that had an associated target B were created. This procedure was applied to compare the dSTORM signal of DG3_63 to mEOS2-DAT, STX1 to DAT, D2R to DAT, PIP2 to DAT, and clathrin to DAT.

#### Quantification of DAT molecules in nanodomains

Primary DA neurons were labeled and imaged for dSTORM with the DAT MAB369 antibody at dilutions 1:200, 1:400, 1:800, 1:1600, 1:3200. Varicosities were manually selected from blinded data files. Clusters of localizations potentially arising from single localizations were identified in the varicosities found in the 1:3200 dilution data set through DBSCAN with the parameters [ε = 30 nm, nps = 2]. A histogram was made comparing the number of localizations found per cluster from this cluster analysis, and 3 localizations was determined to be the estimate for the number of localizations arising from a single monoclonal antibody in this image set. Clusters were then found in the remaining dataset through the same method defined to identify cluster size. A histogram was made to show the number of localizations present within the clusters found with a diameter >75 nm. The peak of this histogram was ∼100 localizations, suggesting there are roughly 33 primary antibodies labeling DAT within a given cluster/nanodomain of this size.

#### Varied label density cluster verification

Varicosities used for the counting experiment were subject to the varied label density cluster verification test (Baumgart *et al*., 2016). Sig for gaussians was set to 15 nm and the cut gauss at x times sig was set to 2. The threshold selected was 2.

#### Labeling and imaging primary cultures with DG3-63

Live primary cultures were labeled with DG3-63 (20 nM) (Guthrie *et al*., 2020) for 10 min in the dark at room temperature and washed 3 times with aCSF (NaCl (120 mM), KCl (5 mM), CaCl_2_ (2 mM), MgCl_2_ (2 mM), 1 mM NaH_2_PO_4_, HEPES (25 mM), glucose (30 mM), pH 7.4). Samples were fixed with paraformaldehyde (3%) for 15 min followed by 3 washes with glycine (20 mM) and NH_4_Cl (50 mM) in PBS. Samples were imaged in normal dSTORM buffer.

#### Transfection of CAD cells, labeling with DG3-63 and pharmacological treatments

CAD cells were transfected with pmEOS2-hDAT encoding mEOS-DAT or with pRC/CMV-hDAT or pRC/CMV-hDAT D421N encoding WT DAT and the DAT mutant D421N, respectively (Herborg *et al*., 2018). The CAD cells (∼1 million cells) were seeded the day before transfection in a 25 cm^2^ flask and washed on the day of transfection before addition of 5 mL of medium. For the transfection, 1 µg of DNA was added to 100 µL of optiMEM as well as 3 µL Lipofectamine (Invitrogen) to a separate 100 µL of optiMEM. After 5 min, the two solutions were mixed and incubated for 30 min before the total mixture was added to the cells. The next day the cells were harvested and seeded on poly-D-Lysine coated coverslips (100,000 cells per coverslip). For DG3-63 labeling experiments, the media was removed the next day from the mEOS-DAT expressing cells and 1 mL of aCSF was added (NaCl (120 mM), KCl (5 mM), CaCl_2_ (2 mM), MgCl_2_ (2 mM), HEPES (25 mM), glucose (30 mM), pH 7.4) before incubation for 10 min with or without 100 µM unlabeled cocaine. Following this, the cells were labeled with 10 nM DG3-63 (Guthrie *et al*., 2020) for 10 min in aCSF. Samples were washed 3 times with aCSF and then fixed for 15 min in paraformaldehyde (3%) followed by 2x wash with glycine (20 mM) and NH_4_Cl (50 mM) in PBS. Samples were imaged in dSTORM buffer as described above. For dSTORM of hDAT WT and D421N, cells the cover slips samples were stained in the same fashion as primary DA cultures. For pharmacology experiments, CAD cells expressing WT DAT were treated with nomifensine (20 µM), noribogaine (20 µM), JHW007 (20 µM) or vehicle for 10 minutes prior to fixation.

#### Pharmacological treatment of acute brain slices

Mice were euthanized with isoflurane and the live brains were removed. Coronal sections, 300 µm thick, were cut with a vibratome in cold aCSF (NaCl (119 mM), KCl (2.5 mM), NaH_2_PO_4_ (24 mM), Glucose (12.5 mM), CaCl_2_ (2 mM), MgCl_2_ (2 mM)) that was being aerated with carbogen gas. Brain slices were transferred to aCSF and maintained for 1 hour at room temperature under constant carbogenation. Slices were then transferred to either aCSF or aCSF with AP5 (100 µM), both at 37° C for 10 min. Slices from normal aCSF were moved to aCSF with or without NMDA (20 µM). The NMDA containing aCSF had reduced magnesium (0.3 mM MgCl2) and added glycine (100 µM). The samples from AP5 were transferred to aCSF with AP5 (100 µM) and NMDA (20µM), reduced magnesium (0.3 mM MgCl_2_), and added glycine (100 µM). These incubated for 5 min, and then the slices were transferred to paraformaldehyde (3%) overnight at 4° C. Slices were transferred to a 30% sucrose solution for 2 days a 4°C and then placed into plastic molds and covered with Tissue-Tek O.C.T. and stored at −80°C. Samples were sliced at 10 µm on the cryostat and stained for dSTORM.

#### Pharmacological treatment of primary cultures

In general, treatments were performed by replacing the media at room temperature with aCSF (NaCl (120 mM), KCl (5 mM), CaCl_2_ (2 mM), MgCl_2_ (2 mM), 1 mM NaH_2_PO_4_, HEPES (25 mM), glucose (30 mM), pH 7.4) followed by stimulation with indicated compounds. For the NMDA or haloperidol experiments with testing of VGCC inhibitors, cells were treated for 10 min with either control aCSF, AP5 (100 µM) or quinpirole (50 µM), ω-conotoxin (1 µM), ω-agatoxin (1 µM). Samples were then treated for 5 min with these same concentrations as cotreatment with either NMDA (20 µM) or haloperidol (10 nM). NMDA treatments were done in aCSF with added glycine (100 µM). All experiments using NMDA were done with aCSF with reduced magnesium (0.3 mM MgCl_2_) for all conditions. For the BAPTA-AM experiment, the pretreatment with BAPTA-AM (25 µM) was for 30 min prior to the 5-min NMDA (20 µM) treatment. For the DAT pharmacology experiments, the neurons were treated for 5 minutes with either nomifensine (10 µM), noribogane (20 µM) or dopamine (200 nM) prior to fixation.

#### SPT-dSTORM

Live primary cultures were labeled with DG3-80 (20 nM) for 10 min and then washed 3 times with aCSF (NaCl (120 mM), KCl (5 mM), CaCl_2_ (2 mM), MgCl_2_ (2 mM – 0.3 mM for NMDA related experiments), 1 mM NaH_2_PO_4_, HEPES (25 mM), glucose (30 mM), pH 7.4). A suitable location to image was found while the sample was kept in aCSF, and then SPTdSTORM imaging was done in a modified dSTORM buffer (560 µg/mL glucose oxidase, 34 µg/mL catalase, 1.4 µL/mL β-Mercaptoethanol in aCSF). Pharmacological treatment were done 5 min before imaging with either 20 µM NMDA with 100 µM glycine, 200 nM haloperidol, or 1 µM UH-232. AP5 (100 µM) was applied for 10 min prior to cotreatment with NMDA. Imaging was performed at room temperature on the NIKON TiE Eclipse TIRF microscope described under dSTORM. Light with a wavelength of 561 nm at 0.3 kW cm^-2^ was utilized to promote blinking of the fluorophores. Images were collected with 16 ms frame rate. Localizations were fit using ThunderStorm as described above. Single molecule trajectories were found using the program swift (Turkowyd et al., 2020). Parameter used beyond the default were: “expdisplacement” : 300,“expnoiserate”: 10,“maxblinkingduration” : 2,“pblink” : 0.5,“preappear” : 0.5,“pswitch” : 0.001,“precision”: 30,“pruningbase” : 2,“pruningrate” : 0.2,“randomseed” : 42,“wdiffusion” : 2,“wdirected” : 1,“wfield” : 0,“wimmobile” : 1.

#### Electrophysiology recordings of DA neurons

DA neuronal cultures were transduced after 3 DIV (days in vitro) with pAAV-pTH-iCre-WPREpA plus either pAAV-hSyn-DIO-mNaChBac-T2A-mKate2-WPREpA, pAAV-hSyn-DIO-mNaChBacMUT-T2A-mKate2-WPREpA or pAAV-hSyn-DIO-Kir2.1-T2A-mKate2-WPREpA. Electrophysiological recording by the whole-cell patch clamp technique were done 2-3 weeks after transduction. The DA neurons expressing either mNaChBac or Kir2.1 were identified based on mKate2 fluorescence using an upright microscope (Olympus BX51WI). For the recordings, the cultures were submerged in circulating, heated, and oxygenated aCSF (in mM; NaCl 125, KCl 2.5, NaHCO_3_ 26, CaCl_2_ 2, MgCl_2_ 1, NaH_2_PO_4_ 1.25, Glucose 25; 2 ml/min, 35°C). The aCSF was supplemented with NBQX (20 µM). The patch glass electrode was filled with (in mM) K-gluconate 122, Na_2_-ATP 5, MgCl_2_ 2.5, CaCl_2_ 0.0003, Mg-Gluconate 5.6, K-Hepes 5, H-Hepes 5, and EGTA 1, and had tip resistance of 4-6 MΩ. After break-in, the cells were held at 0 pA in current clamp mode to measure the pacemaker activity. Recordings were acquired with a Multiclamp 700B amplifier and 1440A Digitizer.

#### Calcium Imaging of DA neurons

Primary DA neurons were prepared as described above but instead of culturing the neurons on glass coverslips, they were cultured on MatTek 35 mm 14 mm glass microwell (no 1.5) glass bottom dishes (MatTek). To enable measurement of Ca^2+^-fluctuations specifically in DA neurons, the neurons were transduced with an AAV encoding double-floxed jRGECO1a (AAV-CAG-Flex-NES-jRGECO1a-WPREpA) together with an AAV encoding Cre recombinase under control of a truncated TH promoter (AAV2-retro-pTH-iCre-WPREpA). The cell medium was changed to aCSF (NaCl (120 mM), KCl (5 mM), CaCl2 (2 mM), MgCl_2_ (2 mM), 1 mM NaH_2_PO_4_, HEPES (25 mM), glucose (30 mM), pH 7.4) immediately prior to imaging. Imaging took place at room temperature on the NIKON TiE Eclipse TIRF microscope described above. Once a suitable neuron was located, the sample was imaged with 561 nm light (10 W cm^−2^). Videos were collected with a 16 ms frame rate. Haloperidol was added mid-video so that the sample had a total concentration of 10 nM Haloperidol. Data was collected from three independent experiments. Two varicosities were identified per image.

#### RNA-seq mining and data analysis

All RNA sequencing data was accessed via the Gene Expression Omnibus (GEO) under the following accession numbers: GSE108020, GSE115070, GSE76381, GSE116470, GSE116138. Raw counts were converted to RPKM and quantile normalized using the preprocessCore (3.11) package in R.

#### Quantification and Statistical Analysis

Statistical details can be found in the figure legends and were computed with GraphPad Prism 8. When employing one-way ANOVA, Dunnett’s post hoc test was used for multiple comparisons. The dSTORM comparisons were normalized to the control sample from the individual imaging session. This was done to control for unavoidable variability in sample preparations and between imaging sessions.

| Experiment | Number of sample preparations | Number of dSTORM Images | Number of varicosities (neuron experiments) or cells (CAD cell experiment) |
| --- | --- | --- | --- |
| IHC dSTORM NMDA | 3 | 102 | 4028 |
| CAD Cell D421N | 3 | 55 | 80 |
| Membrane Potential | 5 | 186 | 849 |
| NMDA and VGCC | 3 | 188 | 841 |
| Haloperidol and VGCC | 3 | 180 | 840 |
| BAPTA-AM | 3 | 95 | 208 |
| STX1 NMDA | 4 | 162 | 678 |
| Clathrin NMDA | 3 | 90 | 304 |
| PIP2 Haloperidol | 3 | 84 | 309 |
| PIP2 NMDA | 3 | 88 | 384 |
| D2 NMDA | 3 | 101 | 344 |
| D2 Haloperidol | 3 | 90 | 524 |
| CAD cell pharmacology | 3 | 179 | 213 |
| Neuron pharmacology | 3 | 237 | 926 |
| Synaptotagmin uptake | 5 | 122 | 893 |

